# Unlocking the Genetic Landscape: Enhanced Insights into Sweet Sorghum Genomes through Comprehensive superTranscriptomic Analysis

**DOI:** 10.1101/2023.09.10.557027

**Authors:** Shinde Nikhil, Habeeb Shaikh Mohideen, NS Raja

## Abstract

Sweet sorghum has gained global significance as a versatile crop for food, fodder, and biofuel. Department of Agriculture, USA declared sorghum a sweet alternative for corn and sugarcane for biofuel production. Its cultivated varieties, along with their wild counterparts, contribute to the core genetic pool. We harnessed 223 publicly available RNA-seq datasets from sweet sorghum to construct the superTranscriptome and analyze gene structure. This approach yielded 45,864 Representative Transcript Assemblies (RTA) that showcased intriguing Presence-Absence Variation (PAV) across 15 existing sorghum genomes, even incorporating one wild progenitor. We identified 301 superTranscripts exclusive to sweet sorghum, encompassing elements such as hexokinases, cytochromes, select lncRNAs, and histones. Moreover, this study enriched sweet sorghum annotations with 2,802 newly identified protein-coding genes, including 559 encoding diverse transcription factors (TFs). This study unveiled 10,059 superTranscripts associated with various non-coding RNAs. The Rio variety displayed elevated expression of light-harvesting complexes (LHCs) and reduced expression of Metallothioneins during internode growth, suggesting the influence of photosynthesis and metal ion transport on sugar accumulation. Intriguingly, specific lncRNAs exhibited significant expression shifts in Rio during internode development, possibly implying their role in sugar accumulation. We validated the superTranscriptome against the Sweet Sorghum Reference Genome (SSRG) using Differential Exon Usage (DEU) and Differential Gene Expression (DGE), which yielded superior estimations. This study underscores the superTranscriptome’s utility in unraveling fundamental sorghum mechanisms, enhancing genome annotations, and offering a potential alternative to the reference genome.

**Significance Statement:** The comprehensive superTranscriptome of seven sweet sorghum genotypes revealed 45,864 genes, including 28.27% novel ones, predominantly comprising non-coding RNAs. Distributing core, dispensable, and cloud genes across 15 sorghum genomes differentiated common genes from cultivar-specific ones. superTranscriptome enhanced the annotation of 14 sorghum genomes with new genes/exons and effectively utilized RNA-seq data to annotate reference genomes. It identified presence/absence variations and non-coding genes and could be a potential alternative to the reference genome.

## Introduction

Sweet sorghum is an ideal crop for future food, feed, and fuel security because of its high cellular biomass, stalk sugar content, and average grain yield. The crop is further characterized by high photosynthetic, water, and nutrient use efficiency; it confers stability under changing environmental conditions (Rao *et al*., 2019). Therefore, it is a potential alternative to fossil fuels to achieve future bioenergy needs. The current breeding schemes need an overhaul to adapt to changing environmental conditions. Instead of focusing solely on recombination within one population, we should consider exploring variability across sorghum species and sub-types. With its diverse primary races and intermediate varieties, Sorghum offers substantial natural variation,(Venkateswaran, Elangovan and Sivaraj, 2018). However, intensive selection and breeding have led to a loss of genetic diversity in modern germplasm, especially in specific agroecological zones (Smith *et al*., 2019).

Past studies suggest that natural selection played an important role in sweet sorghum evolution; non-functional alleles for genes associated with secondary cell wall development qualitatively control midrib color and stem juiciness (Zhang *et al*., 2018). However, the midrib color strongly correlates with traits such as sugar yield, juice volume, and moisture content (Burks *et al*., 2015). Both sweet and grain sorghum show remarkable differences at the phenotypic level, and the genetic makeup of these two crops is also expected to be different. The primary gene pool of sweet sorghum constitutes 35,467 genes along with several Sources of Variation (SV), Copy Number Variation (CNV), and Presence-Absence Variation (PAV); however, deletions are frequent in sweet sorghum, marking an important aspect of sweet sorghum evolution (Cooper *et al*., 2019). The comparative analysis of sweet and grain sorghum genomes observed unique genes associated with variations in sweet sorghum, helping to differentiate these sub-types at the genome level (Zheng *et al*., 2011).

Most genome-wide studies emphasize the number of genes annotated in the reference genome; however, some portion of the genome is only shared by a subset of individuals within the species, termed the dispensable genome (Yao *et al*., 2015). In addition, dispensable genes are associated with complex genomic regions affected by SV and are likely to be missed during the reference genome assembly (Gerdol *et al*., 2020). The comparative studies using reference genomes from a single organism or sub-types are unreliable, as the population shows considerable variation in intraspecies genomes (Bhatti *et al*., 2020). The third revolution in sequencing technologies leads to decreased cost of sequencing (Jiao and Schneeberger, 2017), allowing researchers to sequence more individuals to trap a major portion of dispensable genes among the population. The RNA-seq offers a cost-effective alternative to genome sequencing for identifying functional genes and regulatory elements in plants with complex genomic architecture (Jin *et al*., 2016).

The representative transcript assemblies (RTAs) are a set of genes in the cluster get as a super scaffold and an excellent approach to characterizing core, dispensable, and private genes in a population using RNA-seq datasets (Hirsch *et al*., 2014). In the present investigation, we have constructed RTAs or superTranscripts; these superTranscripts facilitated population-level identification of gene families and non-coding RNAs and helped improve existing sorghum genome annotations.

## Material and Methods

### Sorghum genomes/transcriptomes and pre-processing

The Sweet Sorghum Reference Genome version 2.1 (SSRG), along with 14 published sorghum genomes, including one wild progenitor were retrieved from Phytozome [https://phytozome-next.jgi.doe.gov] and SorghumBase [https://www.sorghumbase.org] from past studies (Cooper *et al*., 2019; Tao *et al*., 2021; Voelker *et al*., 2023). The pan-genome representing 390 diverse bioenergy sorghum accessions was downloaded from the ICRISAT repository [http://dataverse.icrisat.org] (Ruperao *et al*., 2021). The 223 sweet sorghum RNA-seq accessions for seven diverse sweet sorghum genotypes namely Rio (Cooper *et al*., 2019; Y. Li *et al*., 2019), Keller, SIL05 (Mizuno, Kasuga and Kawahigashi, 2016), Della, Dochna (Zhou *et al*., 2022), Roma (Sui *et al*., 2015), and M-81E (Sui *et al*., 2015) from past studies were retrieved from NCBI Sequence Read Archive (SRA) database by using IBM Aspera Connect data transfer protocol and quality trimmed with FastP (Chen *et al*., 2018) to remove poor quality reads and adapter contamination for assembly **(Supplemental Data S1)**. **superTranscriptome construction.** superTranscriptome construction was done by using the Necklace (Davidson and Oshlack, 2018) pipeline with few modifications; it includes individual implementation of genome-guided, *de novo* transcriptome assembly, clustering, superTranscript assembling, and gene expression counts steps for large datasets to avoid time loss due to errors **(Figure 1; Supplemental Figure S1).** The RNA-seq datasets from sweet sorghum cultivars were aligned to SSRG using HISAT2 (Kim *et al*., 2019) with default parameters and known splice site information. The SAM to BAM conversion and BAM sorting were done with SAMtools (Li *et al*., 2009). Genome-guided transcript assemblies for individual samples were obtained by using StringTie (Pertea *et al*., 2015) with default parameters and merged into a single *.gtf file with stringtie---merge option. Further, merged *.gtf was flattened with gtf2flatgtff.pl to extend the gene boundaries for getting longer-length, genome-guided transcripts. Finally, genome-guided superTranscripts were obtained with GffRead (Pertea and Pertea, 2020). The RNA-seq reads further assembled into *de novo* transcripts by using Trinity (Grabherr *et al*., 2011) with--max_memory 100 G. Protein-coding ORFs for *de novo* transcripts were obtained with grain sorghum BTX623, BTX642, and RTX430 proteomes using BLAT (Kent, 2002) with parameters -t = dnax -q = dnax-minScore = 200 and *chimera_braker* tool came along with necklace pipeline. Genome-guided and *de novo* transcriptome assembly transcripts were clustered into groups with parameters-minScore = 200-minIdentity = 90. Finally, a superTranscript representing each cluster/group was constructed using Lace, (Davidson, Hawkins and Oshlack, 2017) with cluster and sequence information. The entire workflow for superTranscriptome construction was divided into three steps and automated with Bpipe, (Sadedin, Pope and Oshlack, 2012). **superTranscriptome quality.** The superTranscriptome assembly quality was analyzed with TransRate (Smith-Unna *et al*., 2016) using sequence information to obtain N50 values. Additionally, superTranscriptome constructed with 98 %, 90%, 80%, and 70% sequence identities were checked with TransRate to see whether clustering influences contiguity. The superTranscriptome and six sorghum genomes were further searched for complete, partial, and missing gene orthologs within the *Poaceae* family with the Poales database using BUSCO (Simão *et al*., 2015). Further, RNA-seq samples from Rio were aligned to superTranscriptome and six published sorghum genomes using HISAT2 (Kim *et al*., 2019) with default parameters to get percentage read alignment and compare RNA-seq read coverage for each assembly.

**Figure 1:**
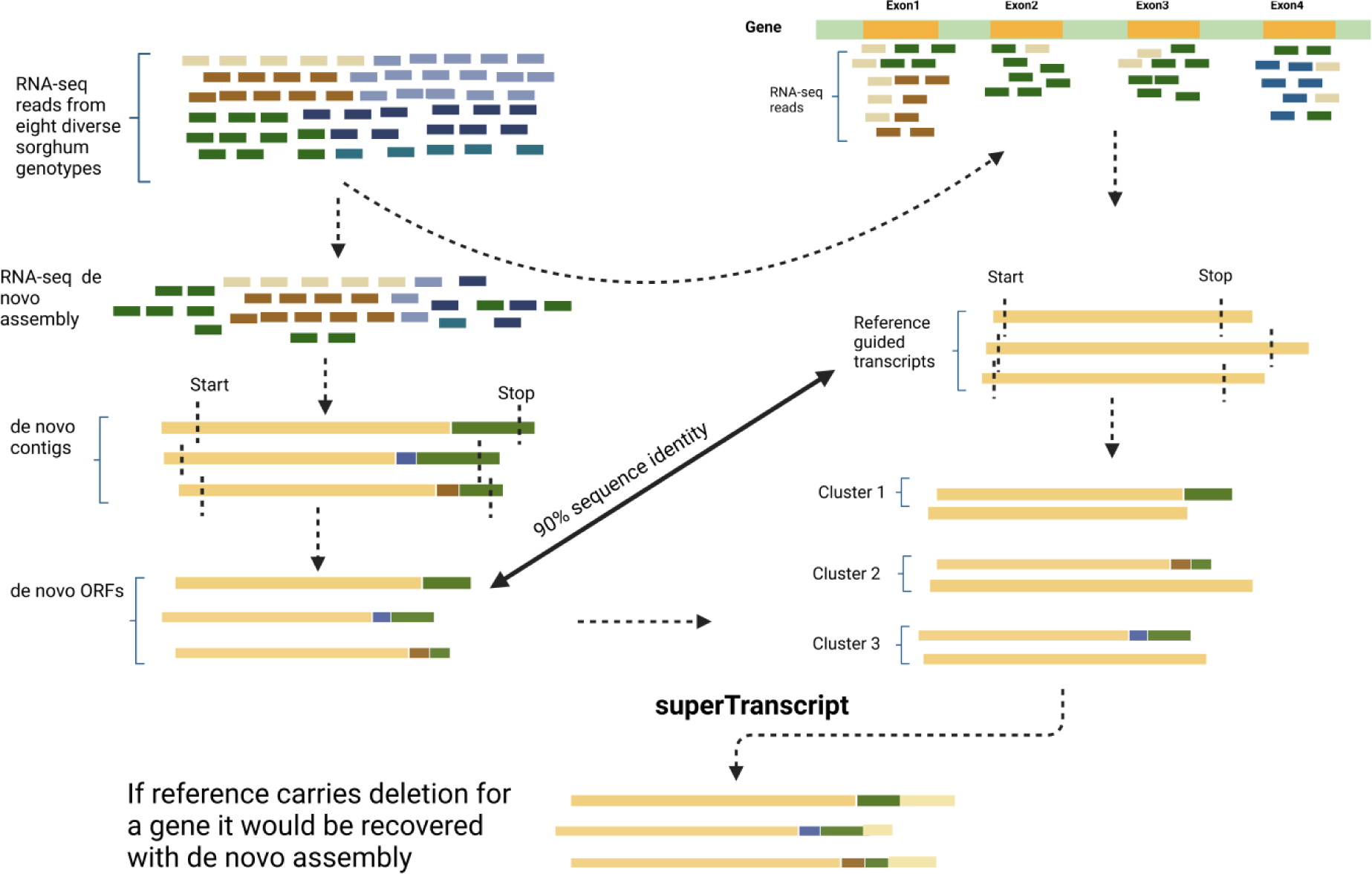
Necklace pipeline (Davidson and Oshlack, 2018) used in present analysis integrates reference guided transcriptome assembly and de novo transcriptome assembly to retrieve complete protein-coding orthologs.

### Functional annotation and characterization of unannotated genes

The genome-guided assembly reported 13815 new genomic regions on SSRG, which were first annotated with SwissProt (Boutet *et al*., 2016) for protein-coding genes using the Trinotate (Ghaffari *et al*., 2014) pipeline; the superTranscripts missing annotations from Swiss-Prot were searched for homology with RNAcentral’s plants non-coding RNA database (Sweeney *et al*., 2020) using BLASTN with parameters pid=90 & qcov=90 for the study of various mi-RNAs and sno-RNAs encoding genes responsible for alternative splicing and silencing. The rest of the superTranscripts that missed annotations from the above two approaches were characterized into putative protein-coding and protein non-coding categories based on their coding potentials with CPC2 (Kang *et al*., 2017) and further annotated with PLncDB; lncRNA database for sorghum by using BLASTN with parameters pid=90 and qcov=90. **superTranscript-based genome annotation and gene densities comparison.** The superTranscripts were used as cDNA/ESTs to annotate SSRG and 14 diverse sorghum genomes using the PASA gene structure annotation pipeline (Haas *et al*., 2008) through the *alignAssembly* method with blat and gmap aligners. superTranscript based annotation comparison of 14 sorghum genomes with existing genome annotation to report novel genes, alternative splice sites addition, and single gene model update using the PASA *annotComapre* method. The alternative splicing updates on 15 sorghum genomes with superTranscriptome using PASA were also done to report transcripts with retained intronic region, spliced exon, and splice sites donors-acceptors. Finally, genome-wide annotation gene structure and slicing updates were visualized as a bar diagram with the *ggplot* R package. The numbers of genes per chromosome for SSRG and superTranscriptome annotated SSRG were obtained, and a bar diagram showing chromosome-wise gene counts was plotted with a custom R script. The superTranscripts were annotated on SSRG using PASA pipeline (Haas *et al*., 2008) produced *.gff3 annotation file and was used to obtain gene densities for tile size 100kb with feature exon by using GFFex function of RIdeogram (Hao *et al*., 2020) R package. Similarly, gene densities for SSRG with past gene annotations were obtained with *.gff3 file from Phytozome from past studies (Cooper *et al*., 2019). Finally, an ideogram highlighting transcribed regions across ten sorghum chromosomes with overlay method was prepared with the same.

### Identification of core, dispensable, and private genes in the population

superTranscripts locations were annotated on 15 sorghum genomes, including wild progenitor 353, using PASA pipeline (F. *et al*., 2006; Haas *et al*., 2008). The superTranscripts shared locations on any genomes with the PASA pipeline were marked as 1 or 0. The core, dispensable, and cloud genes characterization was done based on several genomes’ presence/absence analyses using an approach similar to the previously described (Jobson and Roberts, 2022). Briefly, if superTranscript shared locations on all 15 genomes, it was marked as a core; if they shared locations on 4-14 genomes, they were marked dispensable; and finally, if superTranscripts shared locations on 1-3 genomes were marked as cloud. The superTranscripts exclusively reported on only a single genome were defined as private. The proportion of core, dispensable, and cloud genes per genome was obtained and visualized as a bar diagram, as mentioned in the past investigation on sorghum pan-genome (Tao *et al*., 2021). superTranscripts that did not share any location on the above 15 sorghum genomes were considered orphan genes since they did not share homolog or have partial homolog on the above 15 sorghum genomes (Yao *et al*., 2017).

### Differential exon usage (DEU) analysis

The DEU comparison was performed per the previously mentioned methodology, (Davidson, Hawkins and Oshlack, 2017). The RNA-seq reads of sweet sorghum Rio from past studies (Cooper *et al*., 2019) were aligned to SSRG and superTranscriptome using HISAT2. Novel splicesites were extracted for superTranscriptome to understand blocking within the superTranscripts. The *.gtf file for superTranscript blocks was prepared with the *make_block* tool in the necklace pipeline. The exon bin or blockwise expression count was obtained using FeatureCouts, (Liao, Smyth and Shi, 2014). Finally, statistical significance for exon usage was tested with DEXSeq, (Anders, Reyes and Huber, 2012) R package. Per gene, q-values for differential exon usage were obtained for both with the same. Once per gene q-values were obtained, true positives and true negatives were selected based on q-value cutoffs. The superTranscripts with a q-value below 0.05 were considered true positive, and superTranscripts above a q-value of 0.9 are considered true negative. The Sci-Kit-learn Python module was used to train the datasets with a logistic regression method and report the differences between actual and predicted labels for the above two approaches. The ROC curve showing DEU performance was plotted with a custom R script. Similarly, the confusion matrix for both was also prepared by training datasets with the KNeighborsClassifier method of the Sci-Kit-learn module. Moreover, a DEU comparison between Rio and PR22 during internode development was also performed to report differentially spliced transcripts in these two genotypes.

### Differential gene expression analysis

To report changes in differential gene expression with two different references, i.e., SSRG and superTranscriptome, the RNA-seq reads from leaf, meristem, and internode tissues from six-time points of sweet sorghum Rio were aligned to both, followed by expression counts with FeatureCounts (Liao, Smyth and Shi, 2014) as a transcript count method. Finally, transcript count normalization and log2fold changes were estimated with DESeq2 (Love, Huber and Anders, 2014) R package with Likely hood Ratio Test (LRT) method for time point gene expression normalization. The volcano plots representing DGEs for SSRG and superTranscriptome were plotted using EnhancedVolcano (Blighe K, Rana S and Lewis M, 2022) R package. The top 50 highly expressed genes with P value < 0.05 from the above two methods were visualized as a heat map with the *geom_tile* function of the ggplot2 (Gómez-Rubio, 2017) R package.

Similarly, differential gene expression comparison for Rio and PR22 was done using superTranscriptome as a reference, and the top 50 highly expressed genes (P value < 0.05) were visualized with a heatmap. The gene ontology enrichment analysis was done to identify significantly enriched GO terms between Rio and PR22 internode tissues by preparing a custom GO database with superTranscriptome (additional gene annotations) using AnnotationForge (Pagès *et al*., 2022). Significant GO-enriched terms during internode development in both genotypes were identified using clusterProfiler (Wu *et al*., 2021) R package.

### MSA visualization of agronomically important genes

The agronomically essential genes involved in sugar transport, secondary cell wall development, sugar metabolism, and stress responses were selected for MSA. These include genes encoding SWEET, SUT, INV, Expansin, USP, NAC, MyB, etc. To report new genes identified by superTranscriptome associated with the above gene functions, the superTranscript with missing gene IDs or gene IDs from grain sorghum was retrieved. These are the new gene orthologs in the sweet sorghum population. Finally, multiple sequence alignment of newly reported genes and known genes in sweet sorghum was done using msa (Bodenhofer *et al*., 2015) R package.

## Results

### superTranscriptome construction

Collectively, 223 RNA-seq accessions of seven diverse sweet sorghum genotypes from the public repository formed 45049 genome-guided transcripts with SSRG and 886115 *de novo* transcripts, forming 45864 gene clusters. The representative sequence of each cluster was prepared with Lace, (Davidson, Hawkins and Oshlack, 2017). The superTranscriptome reported 45864 genes with three types of sequences: genome-guided sequences, which are already annotated on SSRG; un-annotated or novel transcribed genomic regions, which require further functional annotation; and *de novo* spliced isoforms or novel genes, which showed less than 90% similarity with SSRG genome-guided contigs but annotated on grain sorghum cultivars genomes such as BTX623, BTX642, and RTX430. The third sequence type is Open Reading Frames (ORFs) from transcripts using protein sequences retrieved from Phytozome (**Figure 2**). With this approach, 886115 *de novo* transcripts were retrieved, but only 163651 were reported with ORFs. Of these 163651, 161712 successfully incorporated known protein-coding superTranscripts from SSRG to retrieve complete protein-coding sequence orthologs. 1939 de novo transcripts did not show significant similarity with any genome-guided transcript formed 815 separate clusters. The superTranscripts number may vary when we change the clustering parameters; it leads to the loss of some annotated genes from the reference genome. However, no significant change in contiguity was reported when we changed the clustering parameters **(Supplemental Table S1)**.

**Figure 2:**
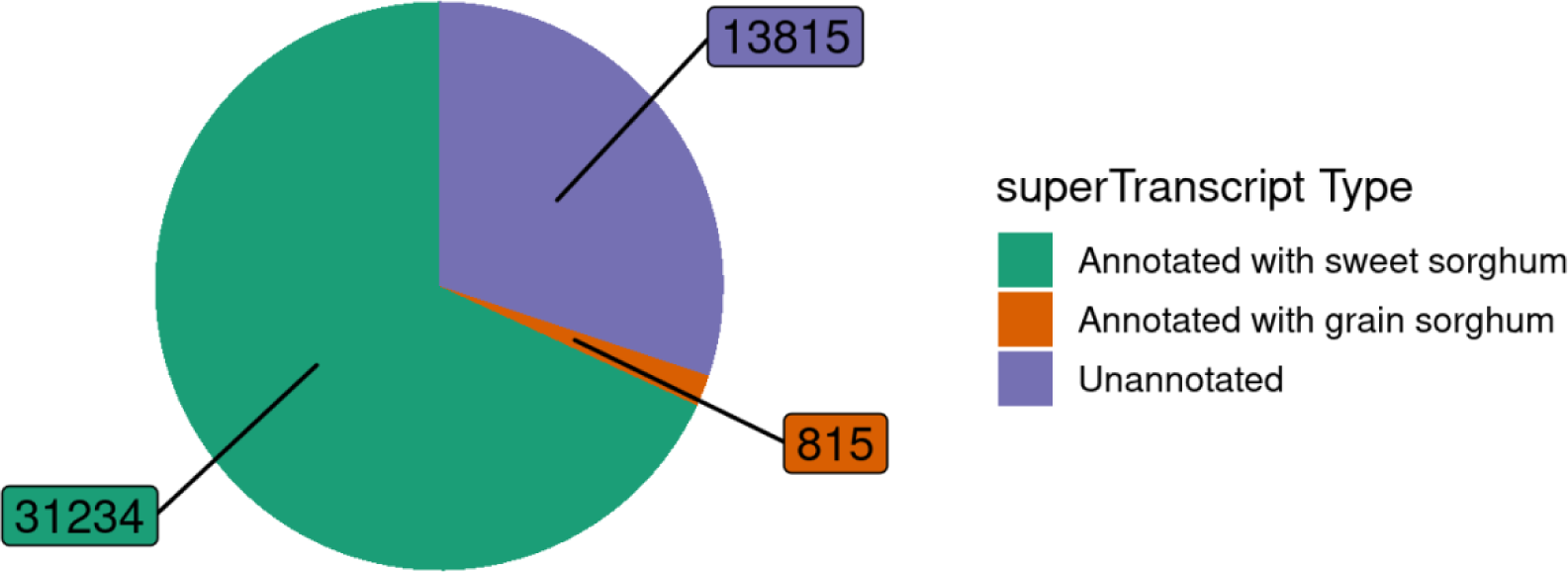
Sweet sorghum superTranscriptome constructed by using necklace pipeline identified with three types of sequences based on their annotations: (1) Annotated with sweet sorghum reference genome (2) Annotated with grain sorghum (3) Unannotated

### superTranscriptome quality

The contiguity analysis of superTranscriptome with the TransRate tool identified the N50 value of the transcripts as 3551 bp. The 30697 (66.93%) and 572 (1.24%) superTranscripts are above 1Kb and 10Kb sequence lengths, respectively. Only 184 (0.4%) superTranscripts are below 200bp sequence length **(Table 1; Supplemental Data S2),** and the rest 14411 (31.42%) sequences have length ranging from 200-800 bp. The result suggests that most sequences are contiguous, could be complete gene orthologs, and highlights the diversity of superTranscript length distribution (**Supplemental Figure S2**). BUSCO assembly completeness analysis suggests that 94.19% of superTranscripts shared complete orthologs with Poales and a 6% duplication rate. Therefore, duplication could be why there are more orthologs for some essential genes (**Figure 3**). The superTranscriptome also reported incomplete/partial genes in comparison with Rio, SC187, BTX642, RTX430, and BTX623 sorghum genomes, suggesting that there are specific genes in the population with missing start/stop codons or both that are playing an essential role in trait development. Assembly quality analysis was done by aligning 48 RNA-seq accessions of sweet sorghum Rio using HISAT2 (Kim *et al*., 2019). The trimmed FastQ reads were aligned to superTranscriptome along with six published sorghum genomes; SSRG showed the highest average read alignment 84.06% among the six genomes, followed by grain sorghum genomes BTX623 (82.29%), BTX642 (82.07%), SC187 (82.06%), RTX430 (81.87%), and superTranscriptome (79.18%) in that particular order; suggest that read coverage is reduced in superTranscriptome over SSRG by 5% (**Supplemental Figure S3)**.

**Figure 3:**
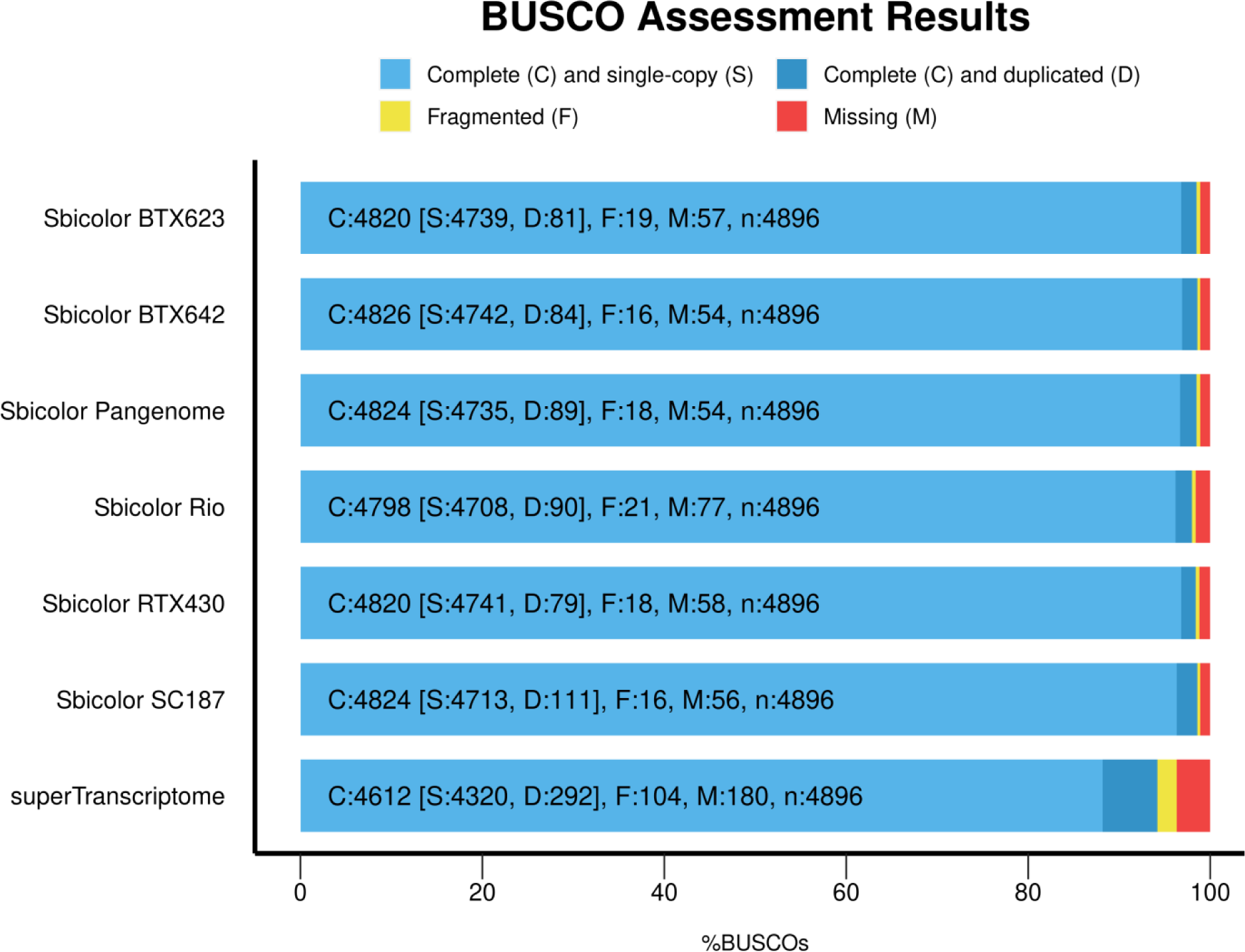
BUSCO assembly completeness analysis with *Poacecae* database by using superTranscriptome and six published sorghum genomes showing complete (single copy/duplicated), fragmented and missing genes.

**Table 1:** Assembly statistics with TransRate for the sweet sorghum superTranscriptome showing number of sequences, total number of bases, the contiguity of the sequences, mean contigs length, smallest and largest sequence in assembly etc.

| Assembly Parameter | Description | Number |
| --- | --- | --- |
| n_seqs | The number of contigs in the assembly | 45864 |
| Smallest | The size of the smallest contig | 96 |
| Largest | The size of the largest contig | 40930 |
| n_bases | The number of bases included in the assembly | 104953927 |
| mean_len | The mean length of the contigs | 2287.73197 |
| n under 200 | The number of contigs shorter than 200 bases | 184 |
| n over 1k | The number of contigs greater than 1,000 bases long | 30697 |
| n over 10k | The number of contigs greater than 10,000 bases long | 572 |
| n with orf | The number of contigs that had an open reading frame | 29610 |
| mean orf percent | For contigs with an ORF, the mean % of the contig covered by the ORF | 47.54242 |
| N90 | The largest contig size at which at least 90% of bases are contained in contigs at least this length | 1205 |
| N70 | The largest contig size at which at least 70% of bases are contained in contigs at least this length | 2405 |
| N50 | The largest contig size at which at least 50% of bases are contained in contigs at least this length | 3551 |
| N30 | The largest contig size at which at least 30% of bases are contained in contigs at least this length | 5065 |
| N10 | The largest contig size at which at least 10% of bases are contained in contigs at least this length | 8726 |
| GC | % of bases that are G or C | 0.48687 |
| bases n | The number of bases that are N | 126 |
| proportion n | The proportion of bases that are N | 0 |

### Functional annotation and characterization of unannotated genes

The sweet sorghum superTranscriptome comprises 45,864 genes. Among these, 31,234 have annotations from SSRG, 815 are from grain sorghum, and 13,815 remain unannotated. Within this unannotated group, the Trinotate pipeline identified and annotated 2,802 new protein-coding genes/superTranscripts with SwissProt. Among these newly annotated protein-coding genes, 559 code for various transcription factors (TFs), including NAC, MYB, and chromo-domain proteins, as determined by their DNA binding domains using PLantTFcat (Dai *et al*., 2013), PLantTFDB (Guo *et al*., 2008) and iTAK (Zheng *et al*., 2016) online server databases. The rest of the 11013 sequences, 572 genes/superTranscripts, were annotated with diverse roles in intron splicing, gene silencing, and ribosomal assembly when searched for non-coding RNA annotation against the RNAcentral database. The remaining 10441 superTranscripts were checked for coding potentials with CPC2 and reported 954 protein-coding and 9487 non-coding regions. Putative 9487 non-coding genes/superTranscripts were potential sources of lncRNAs, of which 6516 were annotated with PLncDB (Jin *et al*., 2021) using BLASTN (**Figure 4; Supplemental Figure S4, Supplemental Data S3**). The results suggest that about 21.93% of sweet sorghum genomes transcribe non-coding RNAs involved in diverse functions, including gene silencing, intron splicing, and gene regulation. Long non-coding RNAs occupied 14% of total expressed sequences; however, this proportion may increase when an independent study is performed.

**Figure 4:**
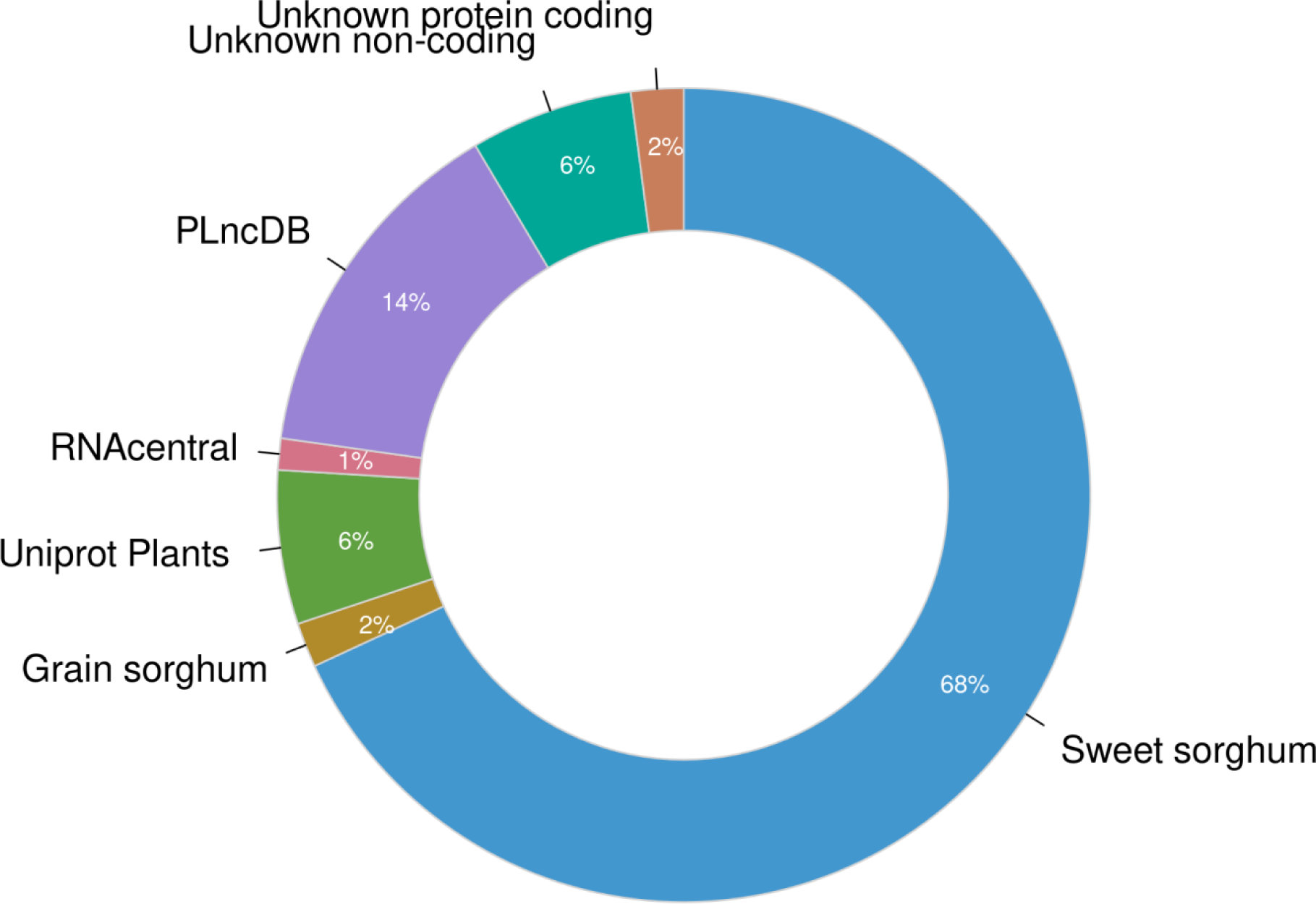
Donut plot showing functional annotations of protein-coding and non-coding sequences in superTranscriptome with different databases.

### superTranscript-based genome annotation

Of a total of 45864, 41169 (89.76%) genes/superTranscripts were annotated on 15 diverse sorghum genomes using the PASA gene structure annotation tool (Haas *et al*., 2008). These genes are selectively transcribed across sorghum genotypes and contribute to variability. The SSRG reported the highest number of annotated genes, 40901 (89.17%), much higher than the previous report, i.e., 35490 in the sweet sorghum (Cooper *et al*., 2019). Additionally, chromosome-wise gene counts and gene densities in SSRG were reported higher when annotated with superTranscriptome using PASA (**Supplemental Figure S5 and S6; Supplemental Data S4)**. The superTranscripts showed several gene structure updates on 14 diverse sorghum genomes (excluding wild progenitor 353). The updates include new gene addition, single gene model update, and alt-splice site addition. Additionally, the superTranscript-based approach reported extensive alternative splicing on these 15 cultivar genomes. The results suggest that superTranscriptome can improve genome annotations, gene structures, alternative splicing, and genome architecture **(Figure 5**; **Table 2).**

**Figure 5:**
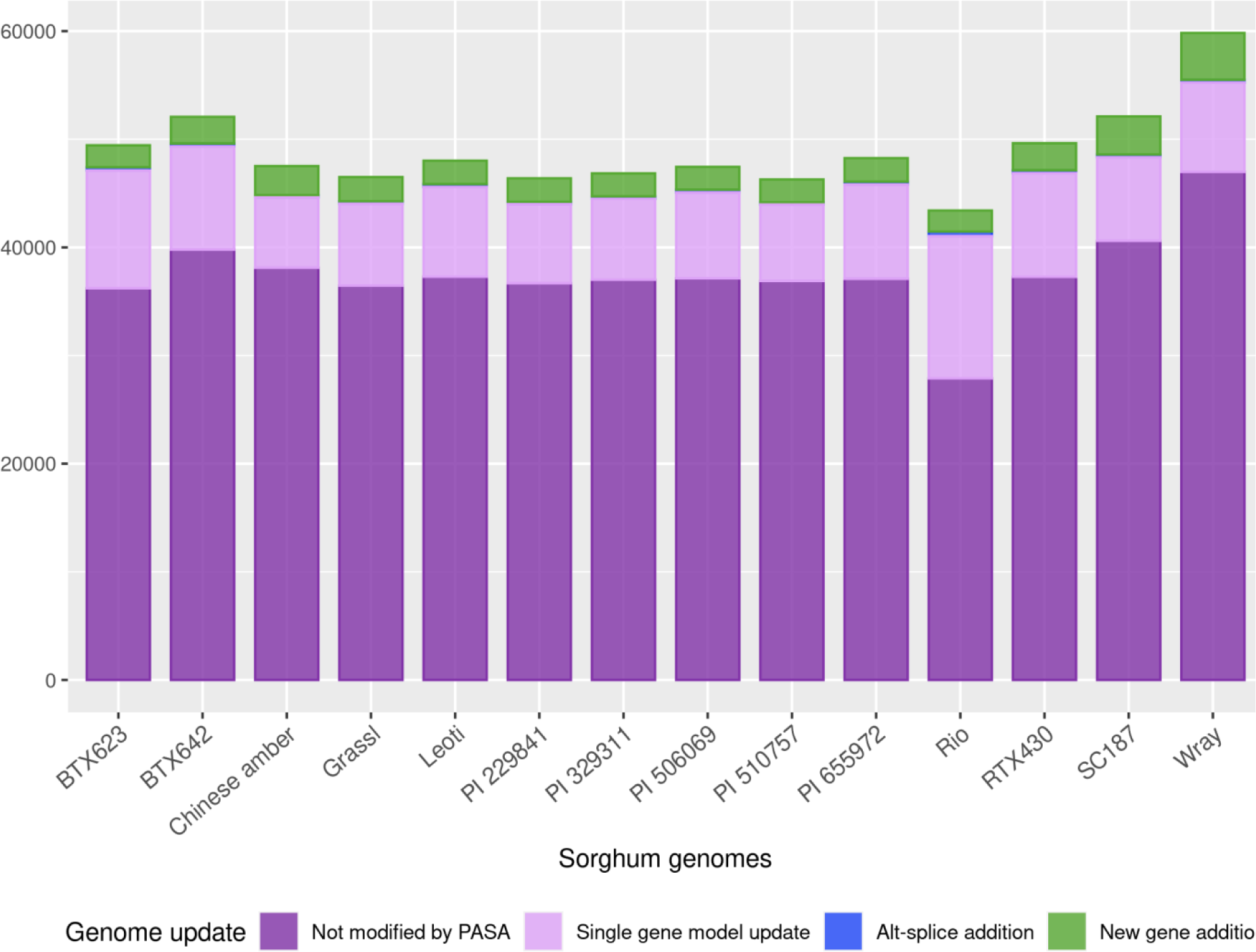
Bar diagram showing various types of structural updates on 15 published sorghum genomes when annotated with superTranscriptome by using PASA gene structure annotation pipeline.

**Table 2:**
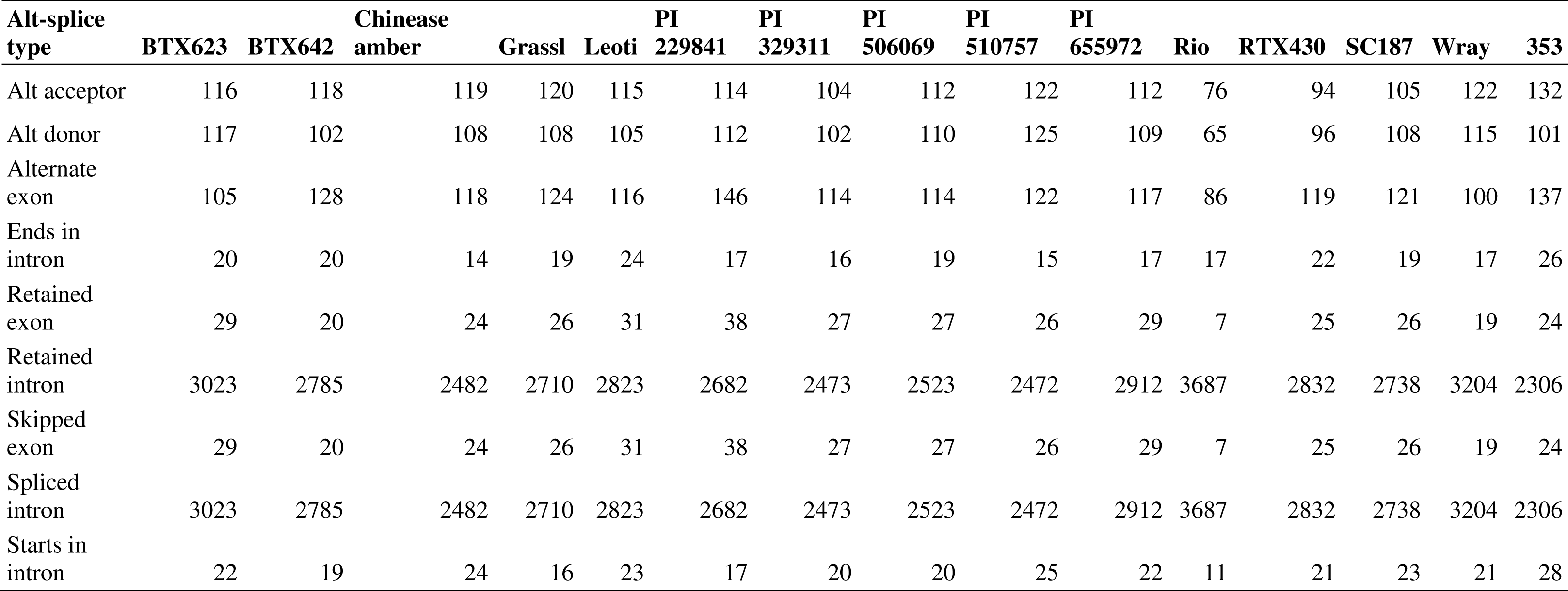
PASA alternative splicing updates on 15 diverse sorghum genomes when annotated by using superTranscriptome; reported with several spliced intron-exons, suggests superTranscripts based approach for genome annotation provides better understanding about gene structures.

### Identification of core, dispensable, and private genes in the sweet sorghum

The Pan-genome analysis classified the gene families into the core, dispensable, and cloud categories for many plants, such as sorghum (Ruperao *et al*., 2021; Tao *et al*., 2021; Wang *et al*., 2021), rice (Sun *et al*., 2017; Zhao *et al*., 2018; Qin *et al*., 2021), maize (Hirsch *et al*., 2014), and pea (Yang *et al*., 2022). The superTranscripts annotated 15 diverse sorghum genomes, including a wild progenitor, based on genomic presence/absence, were classified into 20743 core, 18915 dispensable, and 1511 cloud genes (**Figure 6(a) and 6(b)**). The remaining 4695 superTranscripts were marked as orphan genes because either they lack homologs on the above 15 genomes or have partial homologs on these genomes. Collectively, 24625 (53.69%) superTranscripts were reported with remarkable presence/absence variation on 15 sorghum genomes (excluding single genome superTranscripts and core genes), suggesting that these genes are contributing to variability in sorghum **(Supplemental Data S5, Sheet 2)**. The GO enrichment analysis for dispensable genes suggests that GO associated with Transposable Elements (TEs), proteolytic enzymes, and regulatory elements involved in biological processes showed significant enrichment. This suggests that proteolysis, transposition, and gene regulation of biological processes are significant contributors to the variability observed in sorghum (**Supplemental Figure S7(a))**. Additionally, we reported some superTranscripts that are exclusively located on specific cultivar genomes **(Figure 6(c))**, suggesting that the sweet sorghum carries some genes from cellulosic, grain, and forage backgrounds; however, the function of these genes in sweet sorghum is not known and might carry some modifications. We have reported 301 genes exclusively located on sweet sorghum genomes (Rio, Wray, Leoti, and Chinese amber). These include genes encoding Hexokinase, Core histones (H2A, H2B, H3, and H4), Cytochromes, Glucosyl transferases, Chitinases, cell wall-associated receptor kinases, MYB-TF, and some known/novel lncRNAs. Gene ontology studies of these genes suggest that most genes are associated with various membrane transporters (organic/inorganic ions, electrons, metal ions), catalytic enzymes, cell signaling, gene regulation, chromosomal assembly, and DNA packaging. Sweet sorghum carries additional alleles for cellular homeostasis, energy/carbohydrate metabolism, chromatin regulation, and DNA packaging (**Supplemental Figure S7(b); Supplemental Data S5, Sheet 3)**. Results suggest that these genes are newly evolved in sweet sorghum, exclusively located on sweet sorghum genomes. The orphan genes contribute 9.64% in superTranscriptome, the majority of which codes for TFs, signaling molecules, transporters, and catalytic enzymes that play a significant role in cellular physiology, cell wall development, abiotic and biotic stress resistance and probably help in the sweet sorghum-specific traits when searched for KEGG Orthology (KO) using KAAS (Moriya *et al*., 2007) (**Supplemental Figure S8 (a) and S8 (b)**). No single genome of a particular cultivar of an individual species is enough to cover overall variability in the distribution of genes in that species. Hence, the superTranscriptome could be a suitable alternative for the individual genome to report new genes in the population.

**Figure 6(a).**
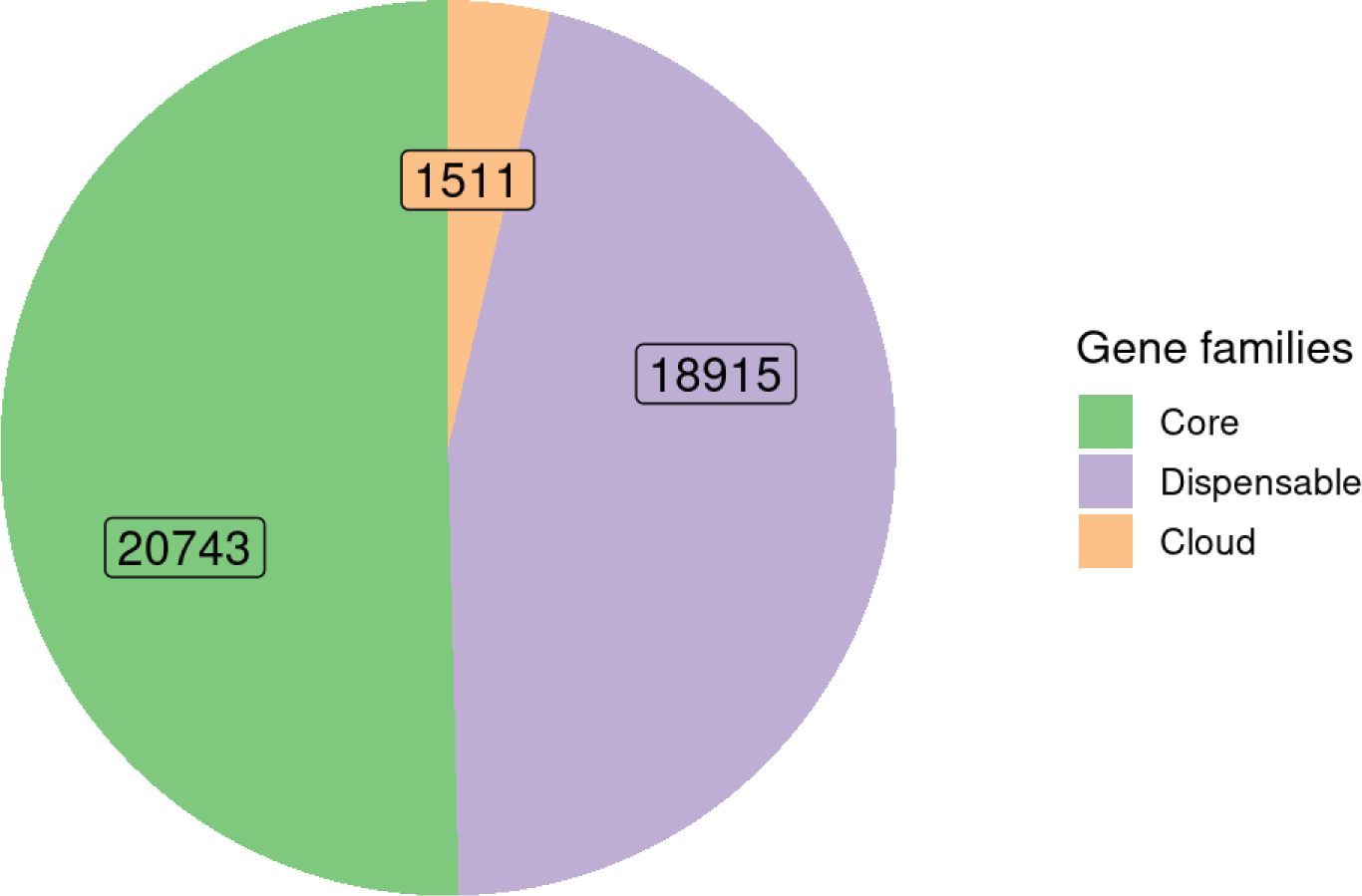
Pie chart showing identification of number of core, dispensable and cloud genes across reported on 15 published sorghum genomes by using superTranscriptome.

**Figure 6(b).**
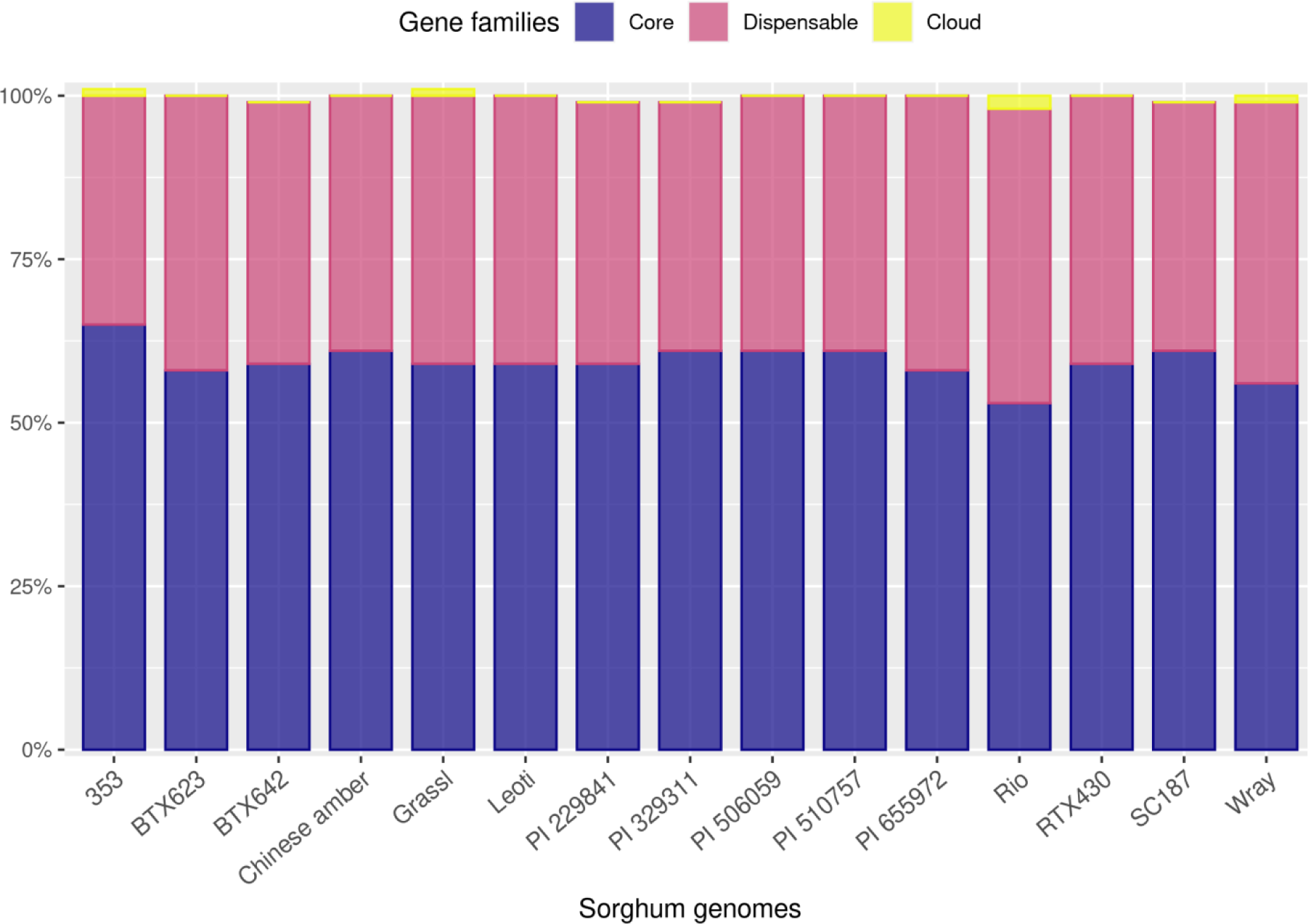
Bar diagram showing proportion of core, dispensable and cloud genes across each sorghum genome when annotated by using superTranscriptome. Flower plot showing distribution of core, shell and private genes across 15 diverse sorghum genomes; helps to report genes from various cultivar background in sweet sorghum.

**Figure 6(c).**
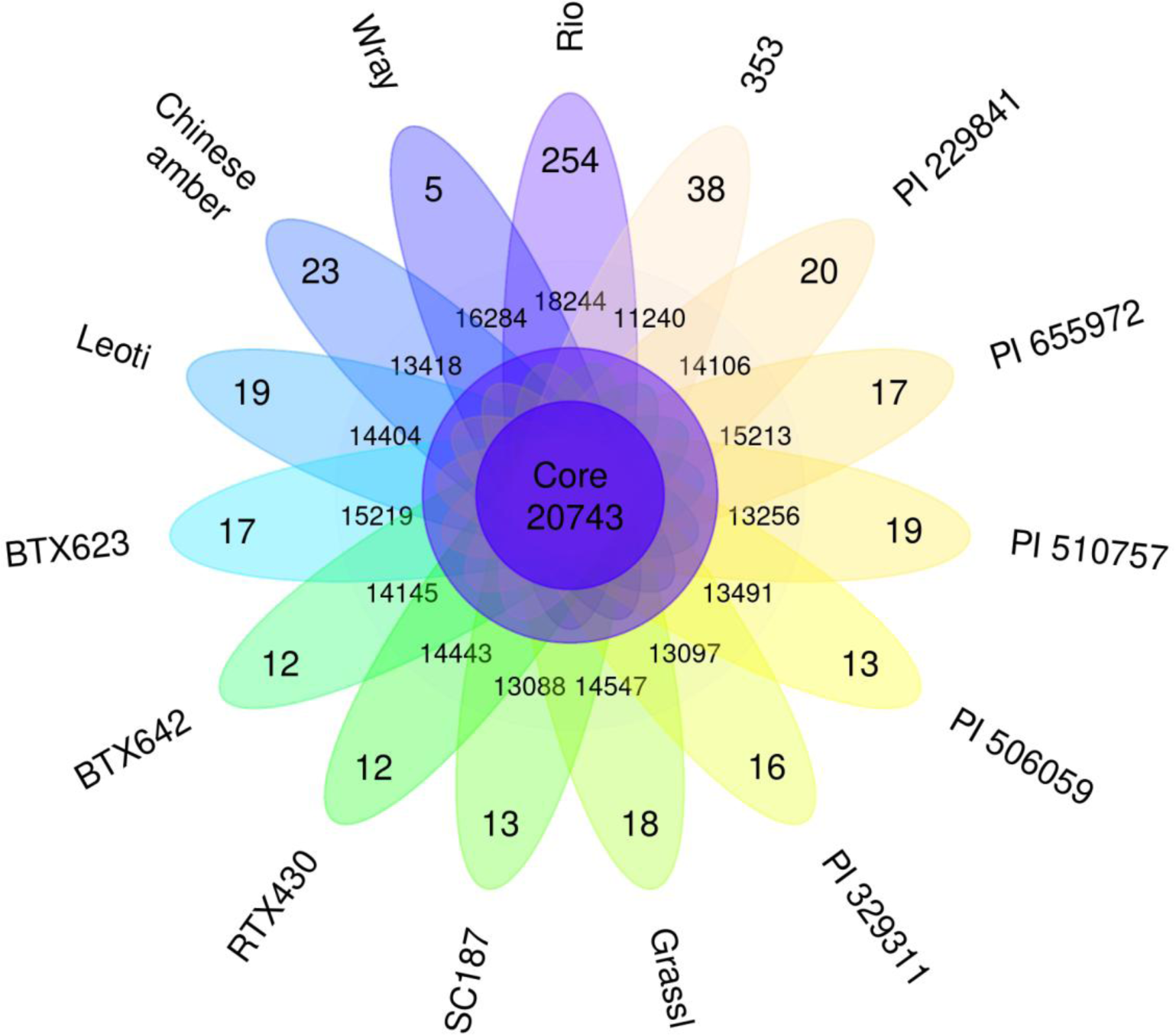
Flower plot showing distribution of core, shell and private genes across 15 diverse sorghum genomes; helps to report genes from various cultivar background in sweet sorghum.

### Differential Exon Usage (DEU) analysis

The traditional genome-based approach with a standard blocking scheme reports only 148847 exons in SSRG. superTranscripts followed a dynamic blocking scheme when aligned with RNA-seq reads and reported with 461355 exon bins, suggesting exon splicing is extensive in superTranscripts. For example, the standard blocking scheme employed for DEU analysis with SSRG yielded five exons for the NLP2 TF coding gene, located on chromosome 6 (SbRio.06G148100), of which three are differentially used for spliced transcript formation under six developmental stages. Where superTranscriptome followed a dynamic blocking scheme for the same gene (NLP2 TF, gene id: SbRio.06G148100) and identified 19 different exon bins, of which eight are differentially used for transcript formation under six developmental stages **(Supplemental Figure S9 (a) and S9 (b); Supplemental Data S6)**. This suggests that superTranscript-based dynamic blocking is more informative for the same gene than the SSRG-based standard blocking. Additionally, this increases the probability of finding more alternatively spliced transcripts; those may be involved in adaptation, stress responses, and trait development. The DEU testing using two references, i.e., SSRG and superTranscriptome, reported 20942 and 22554 genes with differential exon usage. The true positives (with q-value < 0.05) and true negatives (with q-value > 0.9) reported in SSRG and superTranscriptome are 12793 (61.08%), 4583 (21.88%) and 13556 (60.10%), 5749 (25.48%) respectively. The superTranscriptome-based approach found a better classifier for predicted labels than the SSRG one when trained datasets using the logistic regression method **(Figure 7(a))**. Additionally, the confusion matrix prepared with the above two approaches using the KNeighborsClassifier method reported more true labels in superTranscritome than the SSRG-based approach **(Figure 7(b) and 7(c))**. The results demonstrate that the genes identified in the superTranscriptome are indeed true transcripts, as they reported with differential splicing and their ability to provide more accurate estimates of DEU when tested with supervised machine learning. The DEU comparison between Rio and PR22 reported 8787 genes that are differentially used (qvalue < 0.05) by Rio during internode development; however, PR22 has only 5769 (**Supplemental Figure S10**). The GO enrichment analysis of these genes identified that most of them are involved in cellular transport, cellular trafficking, photosynthesis, protein folding, nucleotide metabolism, post-transcriptional and translational modification, etc., suggesting that differential splicing of these genes during internode development could be the reason for sugary internodes **(Supplemental Figure S11)**.

**Figure 7(a).**
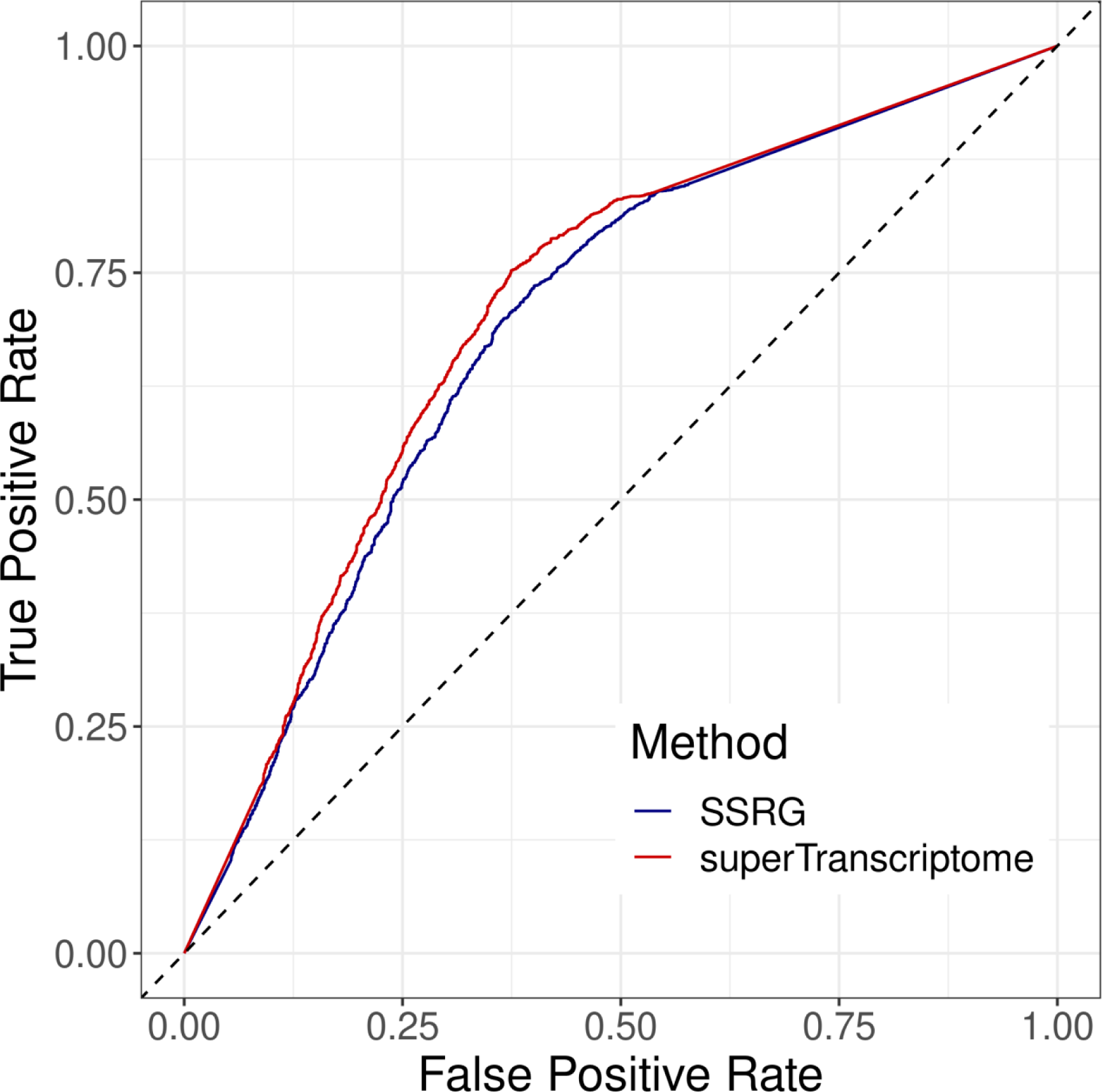
ROC curve based on TPR and FPR values with logistic regression method showing superTranscritptome with dynamic blocking giving better estimates of DEU than SSRG based standard blocking approach.

**Figure 7(b).**
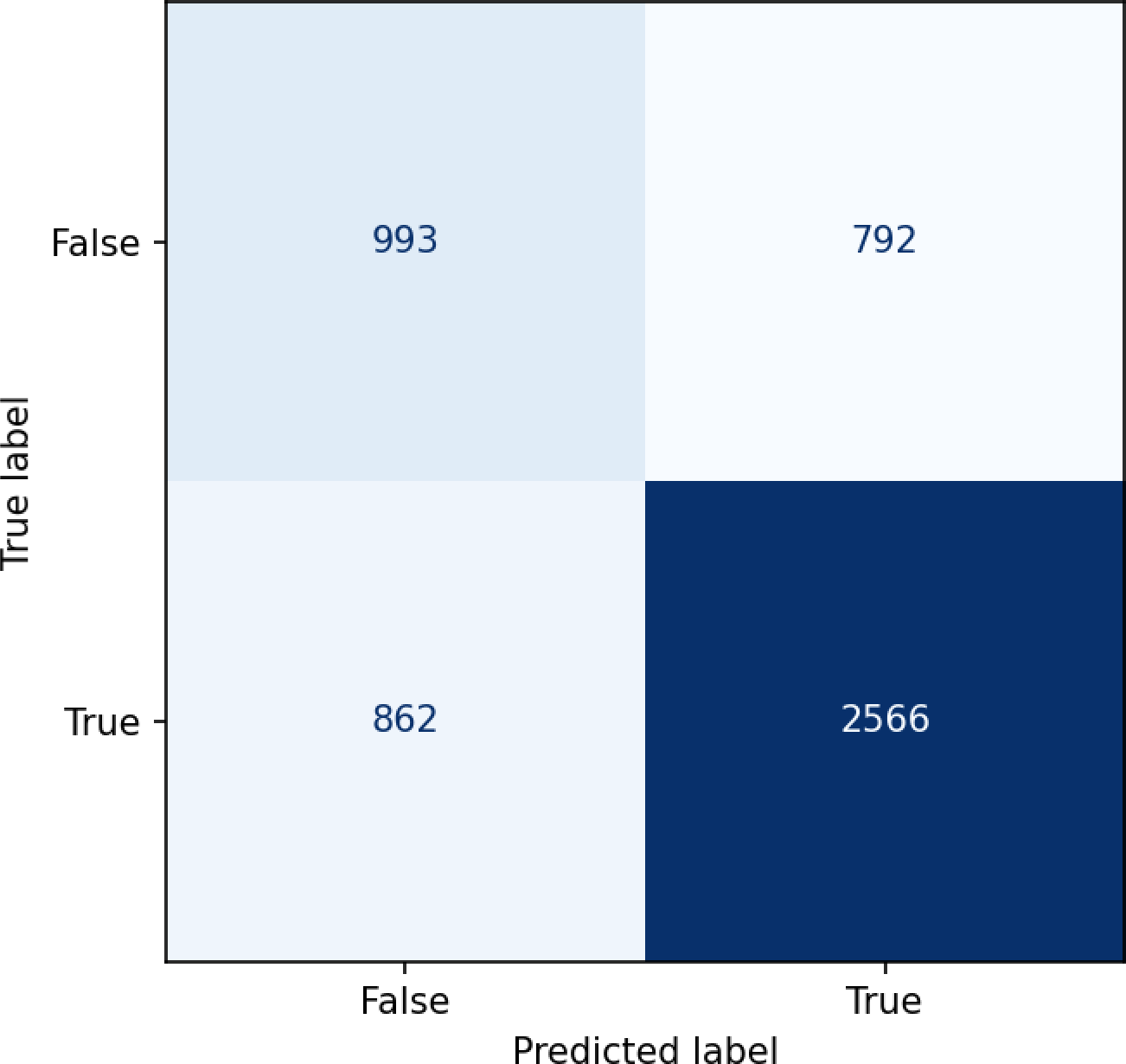
Confusion matrix prepared for DEU with SSRG showing differences between true and predicted labels with 70% train and 30% test datasets by using KNeighborsClassifier method.

**Figure 7(c).**
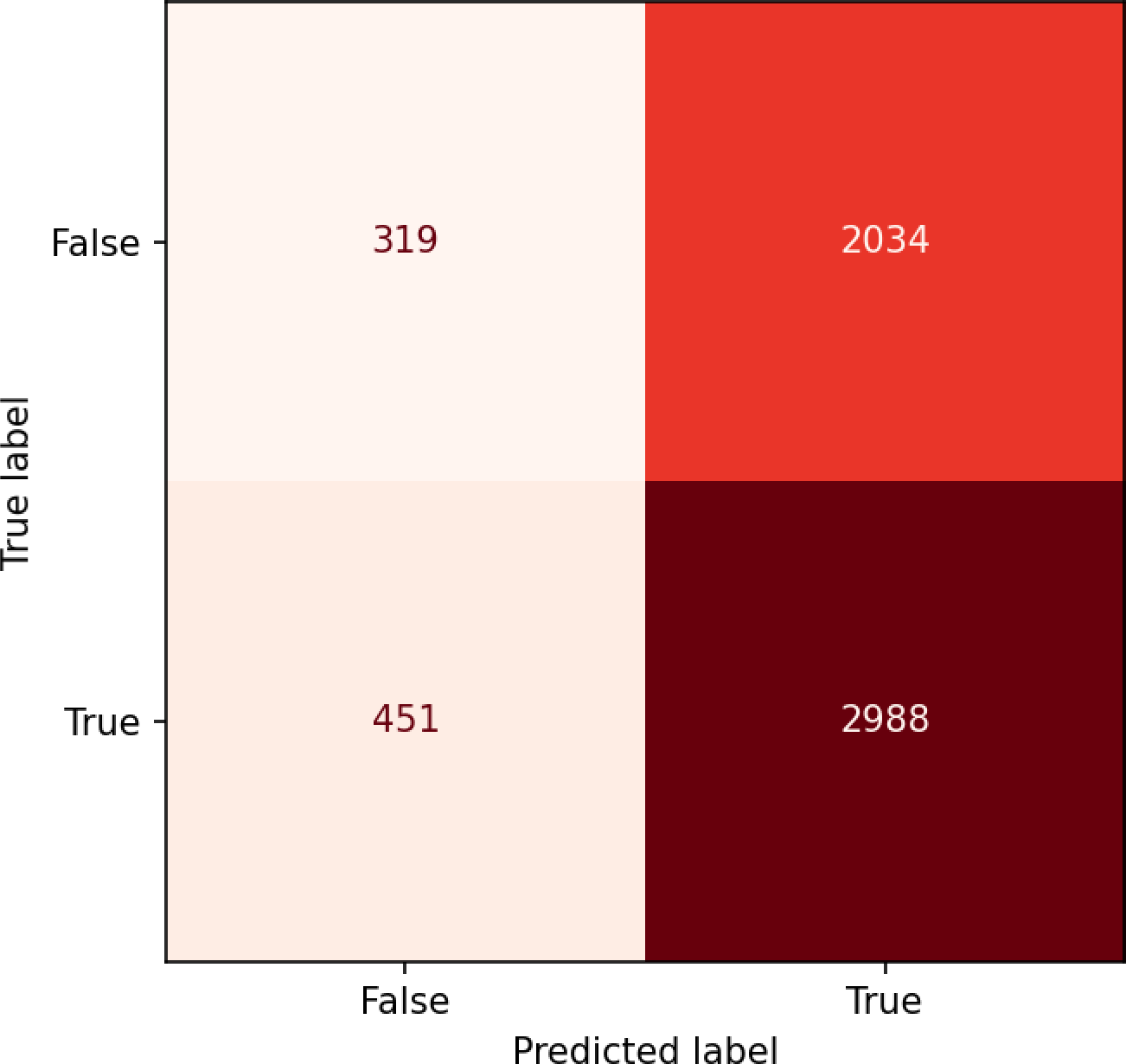
Confusion matrix prepared for DEU with superTranscriptme showing differences between true and predicted labels with 70% train and 30% test datasets KNeighborsClassifier method, suggests number of true labels predicted are higher in superTranscriptome.

### Differential gene expression (DGE) analysis

Gene-level abundance estimates and DGE analysis using SSRG and superTranscriptome as a reference identified a total of 20940 and 22036 genes, respectively, and showed significant (P-value < 0.05) changes in gene expression across the leaf, meristem, and internode tissues over six-time points (**Figure 8(a) and 8 (b); Table 3; Supplemental Figure S12 (a))**. Among 50 highly expressed genes in the leaf, meristem, and internode tissues, 83% are the same with the above two references found, and the rest of the 17% genes showed slightly altered gene expression (**Supplemental Figure S12 (b) and S12 (c)**). Our analysis of the ROC curve prepared by training datasets (70% train and 30% test) with logistic regression method reported superTranscriptome gives better DEU estimates than SSRG **(Supplemental Figure S13).** superTranscript-based approach identifies more truly expressed genes than SSRG and highlights the suitability of superTranscriptome for DGE estimates. Past studies with reference genome and *de novo* transcriptome-based DGE analysis showed slight variation in the gene expression levels (Davidson and Oshlack, 2014). In our study, the DGE comparison between Rio and PR22 during internode development reports several Light Harvesting Complexes (LHCs) that are highly expressed in Rio than in PR22 and shows that active photosynthesis is helping in sweet phenotype development. Interestingly, Rio was found with reduced expression of Metallothionein-II during internode development **(Figure 8(c))**, but in meristem tissues, higher levels of expression (P-value < 0.05). It suggests that metal ion transport is selectively controlled among tissue types and plays a vital role during vegetative growth or mitosis. However, metal ion transport is reduced after the cell matures, which confers a sweet phenotype in Rio. Differential gene expression reported GO terms associated with metal ion transport, secondary metabolite transport, cellular trafficking, etc., are enriched in PR22 during internode development **(Supplemental Figure S14)**. Therefore, active transport of metal ions, secondary metabolites, and organic compounds contributes to dry or pithy stems in sorghum. Further, the genes reported with significant (P-value < 0.5) changes in expression during internode development in both genotypes have 9028 common, 6092 unique to Rio, and 2840 unique to PR22, respectively. **(Supplemental Figure S15)**. Interestingly, 332 and 193 lncRNAs were reported exclusively to Rio and PR22, respectively, with significant (P-value < 0.05) changes in expression during internode development, suggesting that lncRNAs could be hidden players that control internode development and confers sugary internode **(Supplemental Figure S16 (a) and S16 (b))**.

**Figure 8(a):**
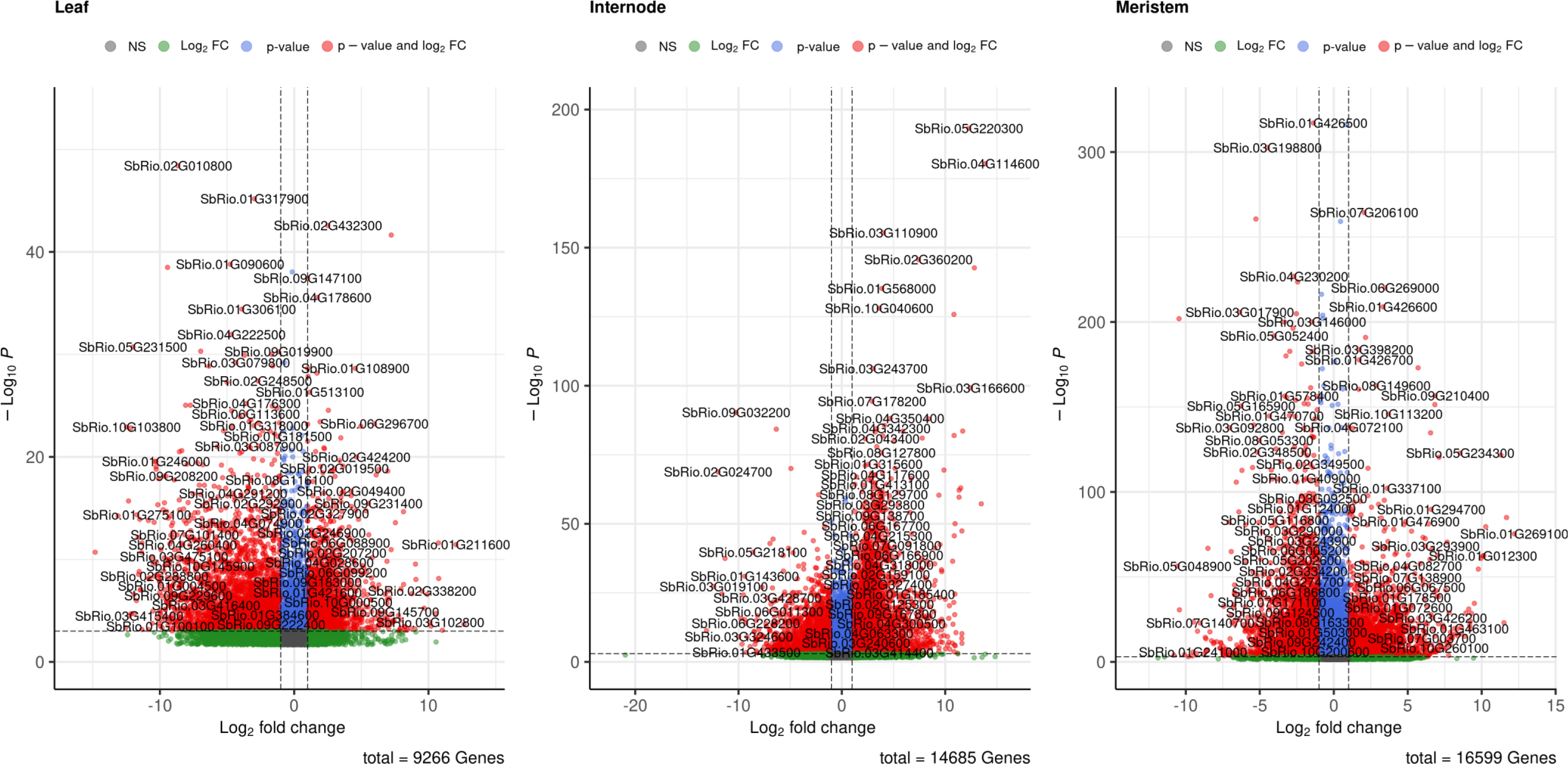
Volcano plot showing genes with drastic changes in expression across Leaf, Internodes and Meristems tissues when used SSRG as a reference for DGE analysis.

**Figure 8(b):**
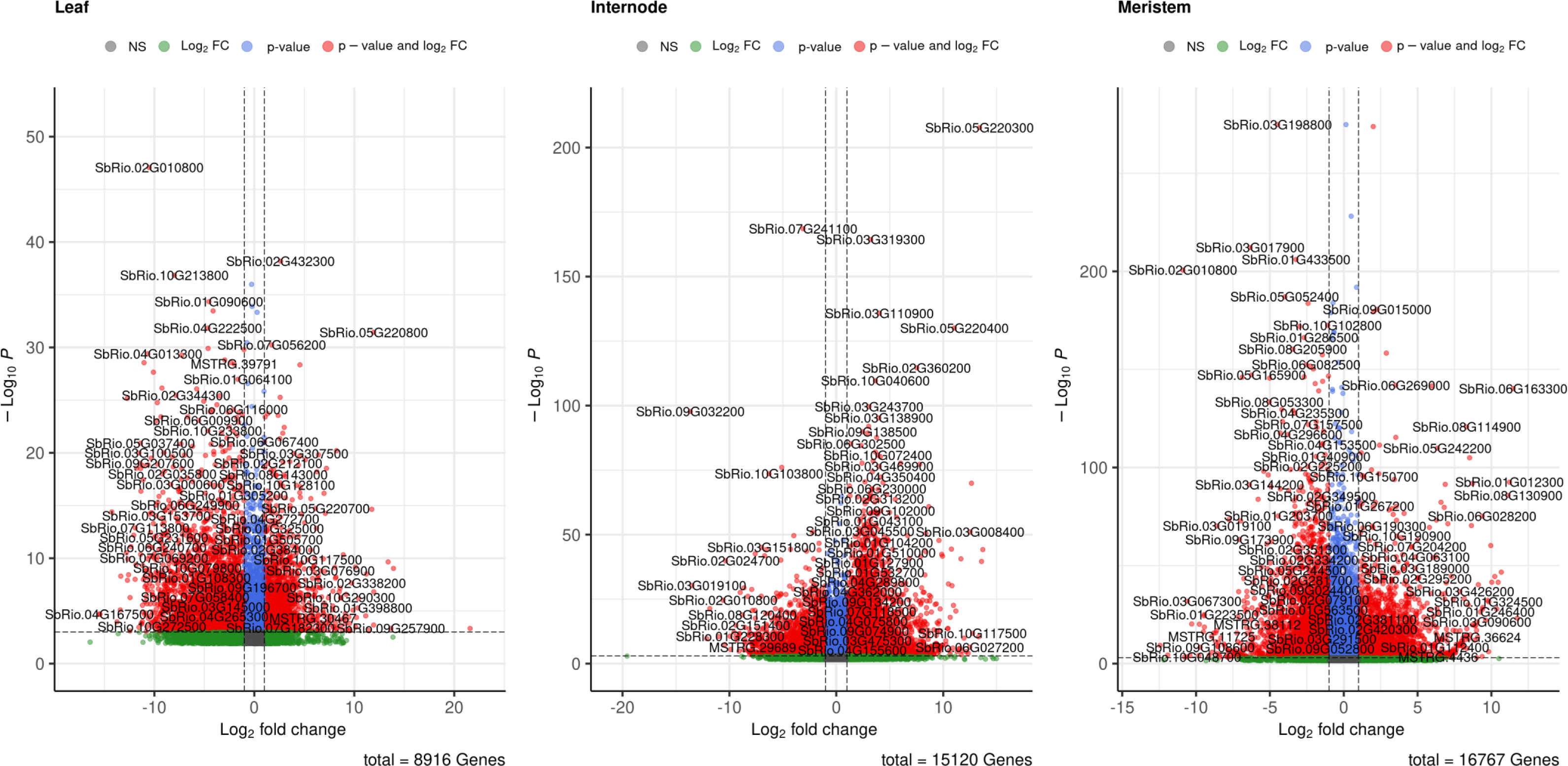
Volcano plot showing genes with changes in expression across Leaf, Internodes and Meristems tissues when used SSRG as a reference for DGE analysis, suggest that more number of genes are reported significant with changes in expression with superTranscriptome based reference.

**Figure 8(c):**
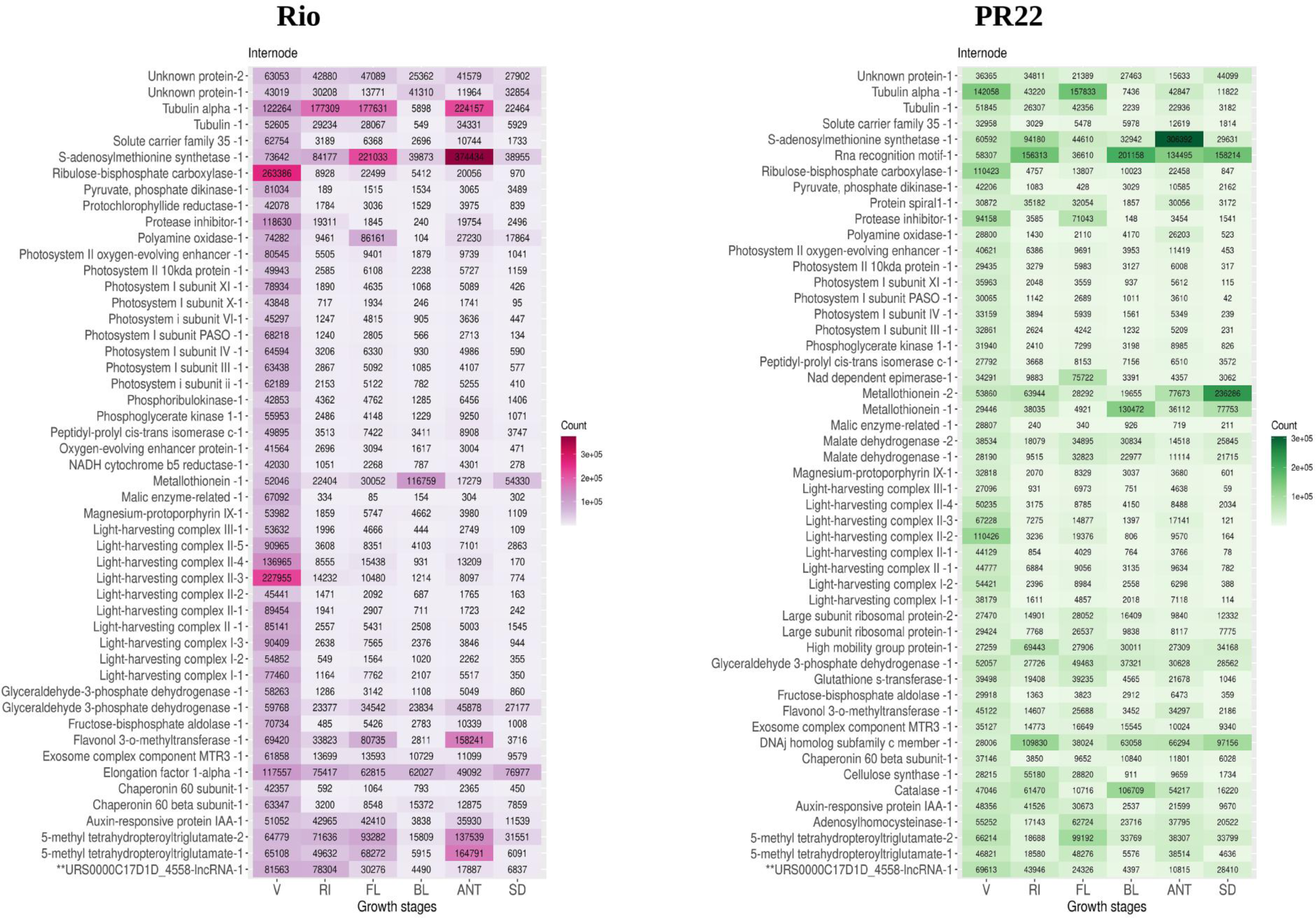
Heatmap showing Top 50 highly expressed genes in internode tissues of Rio and PR22, reports more LHCs are expressed in Rio along with missing expression of *Metallothionein II* during internode development.

**Table 3:**
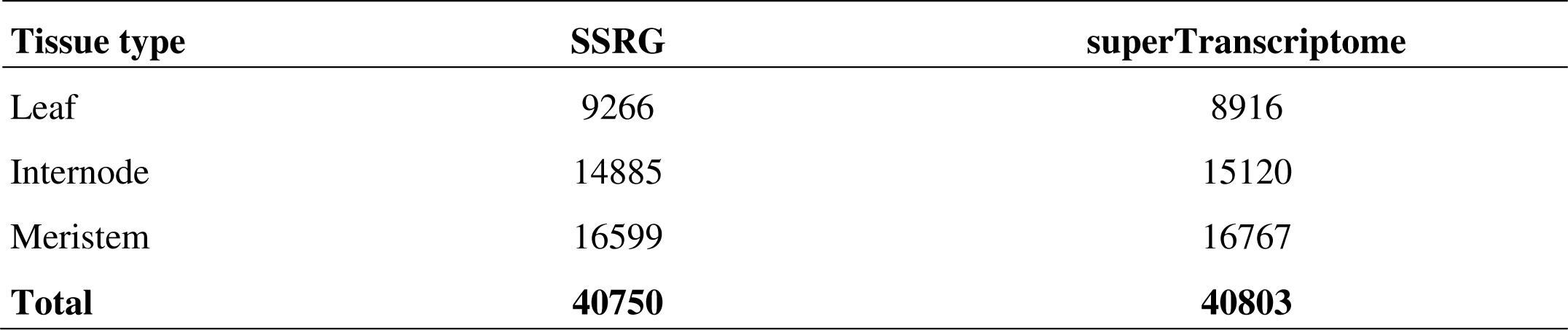
Differential Gene Expression (DGE) analysis performed using DESeq2 with two different references reported number of genes showed significant (P value < 0.05) changes in expression across leaf, meristem and internode tissues of sweet sorghum Rio.

### Identification of agronomically essential genes and MSA visualization

The agronomically important genes such as SWEET, SUT, INV, Expansin, USP, NAC, and MYB reported copy numbers in superTranscriptome than SSRG. The SUT4 gene is one of the six SUT genes identified in sweet sorghum; however, this gene is missing a location in SSRG due to large deletions (Cooper *et al*., 2019). The present analysis reported a SUT4 ortholog from *de novo* assembly. The Sugars Will Eventually Exported Types of Transporters (SWEET) are gene families involved in phloem loading and unloading. The superTranscriptome approach reported 24 SWEET genes with one additional SWEET locus, i.e., SWEET13 located on chromosome 8 with only one exon. A result suggests that the newly reported SWEET gene is also partial in sweet sorghum. The plant-specific NAC transcription factor qualitatively controls dry biomass, sugar production, and grain yield (Xia *et al*., 2018). superTranscriptome identified 130 NAC-TF encoding loci, of which six are newly reported. The number of NAC genes identified in the superTranscriptome is comparatively less than previously reported in the sorghum genome (Kadier *et al*., 2017) and loss of NAC genes might be the reason for sweet phenotype development. Although we retrieved more and complete gene orthologs for agronomically important genes by integrating *de novo* transcriptome assembly and genome-guided assembly approaches, the functionality of newly reported SWEET, SUT, and NAC-TF in sweet sorghum could not be established because of the presence of uni and di exonic loci. Further analysis, such as GWAS, may provide more information about these genes. The agronomically important genes were subjected to MSA analysis followed by phylogram construction, showing a shared phylogenetic relationship with existing genes in the genome (**Supplemental Figure S17 – S22**).

## Discussion

superTranscriptome identified a total of 45864 genes, which is higher than the previously reported in sorghum (McCormick *et al*., 2018), sweet sorghum (Cooper *et al*., 2019), sorghum pan-genome (Ruperao *et al*., 2021; Tao *et al*., 2021; Wang *et al*., 2021), rice pan-genome (Zhao *et al*., 2018; Qin *et al*., 2021), barley (Jayakodi *et al*., 2020), and much lesser than maize pan-genome (Hufford *et al*., 2021). The results validate that the superTranscriptome approach identified more expressed genes and suggests that a single reference genome cannot report all genes expressed in sweet sorghum. Since superTranscripts are probabilistic gene models for sweet sorghum, their functionality may vary from genome to genome depending on the gene structure. The N50 value for superTranscriptome is 3551 bp, much higher than the N50 value previously reported for pan-transcriptome assemblies of maize (Hirsch *et al*., 2014; Jin *et al*., 2016), alfalfa (Medina, Samac and Yu, 2021), potato (Petek *et al*., 2020), and barley (Ma *et al*., 2019), showed the increased contiguity in superTranscriptome assembly. BUSCO assembly completeness studies reported gene duplication and missing/partial genes that could be the reason for trait development in sorghum. Also, past studies on sorghum pan-genome reported BUSCO gene completeness ranging from 94.4 to 98% in the sorghum (Tao *et al*., 2021), supporting our findings. Past studies reported partial genes were associated with the sugarcane aphid (SCA) resistance for the cultivated sorghum variety TX278 (Wang *et al*., 2021). Furthermore, a comparative analysis of maize genomes also reported that more incomplete genes and missing genes contribute to maize adaptation under diverse environments and crop improvement (Yang *et al*., 2017; C. Li *et al*., 2019). The superTranscriptome reported an average of 5% less read coverage over SSRG. We have lost an average of 5% transcribed sequences over SSRG during superTranscripts construction or duplicated genes caused reduced alignment due to multi-mapping.

The lncRNAs contribute 14% of superTranscriptome. However, past studies on lncRNAs suggest they are highly tissue-specific and condition-specific (Statello *et al*., 2021). In the present analysis, we reported significant changes in expression during the internode development. The total number of lncRNAs identified in maize (Wang *et al*., 2015; Lv *et al*., 2019), sorghum (Sun *et al*., 2020), rice (Zhou *et al*., 2021), and barley (Unver and Tombuloglu, 2020) ranged from 8000 to 23309 and showed that lncRNAs occupy the majority of total RNA. Recent studies on ten high-quality genome assemblies of diverse bioenergy sorghum genotypes reported similar results for lncRNAs (Voelker *et al*., 2023). The dispensable genes are major players contributing to variability that codes for various Transposable Elements (TEs), proteolytic enzymes, and regulators of biological processes; its proportion is about 53.69 % in the superTranscriptome. However, previous studies reported 63.6% dispensable genes in sorghum when compared with rice, soybean, and *Brachypodium sp*., suggesting that sorghum is more genetically diverse than other crops (Xin *et al*., 2021). The past studies on sorghum reported that Transposable Elements (TEs) may play a role in the gene content variation (Voelker *et al*., 2023) and our work substantiated it but processes, such as proteolysis and gene regulation, were not reported. Therefore, proteolysis and gene regulation also contribute to the observed sorghum variability. We identified 301 genes exclusively located on sweet sorghum genomes, including hexokinases, core histones, cytochromes, etc. These genes may be newly evolved in sweet sorghum. A recent study on sorghum reported that some histones and unique chromatin remodeling factors were newly evolved (Hu *et al*., 2022).

Our DEU analysis reported more true positive genes in the superTranscriptome-based approach than SSRG when datasets were trained with the KNeighborsClassifier method and gave better estimates of DEU when trained with the logistic regression method. superTranscriptome reported more differentially spliced genes; therefore, they are true transcripts. Additionally, the count-based methods we have used for gene-level abundance estimates and statistical inferences are superior to traditional TPM-based methods (Soneson *et al*., 2016). DEU comparison between Rio and PR22 during internode development suggests that differential splicing is extensive in Rio and could be the reason for the sugary internode.

The DGE, by using two different references, i.e., superTranscriptome and SSRG, reported more DEGs (P-value < 0.05) with superTranscriptome than SSRG. superTranscriptome gives better estimates of DGE than SSRG when trained datasets with the logistic regression method. DGE comparison between Rio and PR22 during internode development reported more LHCs and missing expression of Metallothionein-II in Rio under the Top 50 category, suggesting that higher expression of LHCs and lower expression of Metallothioneins contributes to sugary internode. Metallothioneins are active transporters of various metal ions, including copper (Cu^2+)^, and deficiency impacts Cu^2+^ accumulation in various plant parts (R. Benatti *et al*., 2014). The past studies on *Chenopodium murale* (Llerena *et al*., 2021)*, Colobanthus quitensis* (Contreras *et al*., 2018), and sugarcane (Agarwala *et al*., 1993) reported copper concentrations stimulate sugar levels in various plant parts, indicates the presence of the metal ions inside the cell positively regulates sugar accumulation in internode tissues.

SUT, SWEET, and NAC2–TF are important gene families controlling internode development in sorghum (Mizuno, Kasuga and Kawahigashi, 2016; Zhang *et al*., 2018). Past studies reported six SUT genes in sorghum with abundant gene expression in sweet sorghum stem internodes compared to grain sorghum (Li *et al*., 2014; Babst *et al*., 2021). The SUT4 is reported with putative deletions in Rio (Cooper *et al*., 2019); our *de novo* transcriptome assembly analysis with 223 RNA-seq accessions reported its presence, suggesting that SUT4 is present in the sweet sorghum population. Several SUT genes reported in maize (4-7) (Leach *et al*., 2017), rice (5)(Aoki *et al*., 2003; Hirose *et al*., 2010), wheat (4)(Deol *et al*., 2013), and barley (5)(Radchuk *et al*., 2017) with a variety of functions in growth and development. To date, 23 SWEET genes have been reported in the sweet sorghum (Mizuno, Kasuga and Kawahigashi, 2016); however, SSRG reported only 21 SWEET genes, along with two with putative deletions, namely SWEET3–3 and SWEET8–2); but we have reported 24 orthologs using a superTranscriptome-based approach. Interestingly, maize and foxtail millet also have the same number (24) of SWEET orthologs, (Liu, Fan and Ma, 2022), suggesting that there could be a history of introgression of SWEET genes between sorghum, maize, and foxtail millets. Genome introgression between wild relatives and crops within the same family was reported in other crops also (Ellstrand, Prentice and Hancock, 1999; Hufford *et al*., 2013; Ananda *et al*., 2020). In sorghum, plant-specific NAC-TFs control a verity of traits, including high cellular biomass (Xia *et al*., 2018), insect-pest resistance (Zhang *et al*., 2013), and drought/salinity tolerance (Sanjari *et al*., 2019; Punia *et al*., 2021), etc. Natural mutants for the dry gene on chromosome 6 contribute to sugary phenotypes in sorghum (Zhang *et al*., 2018). In this study, we have reported six new loci for NAC-TF, one with secondary cell wall development, i.e., NAC-73-2* with remarkable PAV on 15 genotypes, proposing that this gene contributes to variability in sweet sorghum.

### Conclusion

Sweet sorghum superTranscriptome reported with 45864 genes, of which 301 are exclusively located on sweet sorghum genomes and controlling a variety of functions, including chromatin organization, gene regulation, sugar metabolism, and cell wall synthesis; these genes might have newly evolved in sweet sorghum. The orphan genes reported in sweet sorghum codes for various TFs, signaling molecules, and catalytic enzymes confer additional traits such as resistance. Our DEU and DGE analysis suggest that Rio has more differentially spliced genes than differentially expressed genes, suggesting differential splicing might lead to a sugary internode. Interestingly, Rio reported more LHCs with elevated expressions in Leaf, Meristems, and Internodes, highlighting their increased photosynthetic efficiency and lower expression of metallothioneins. This indicates that the metal ion transport and photosynthesis controls sugary internode formation. The superTranscriptome reported a 6% duplication rate with BUSCO analysis, leading to more orthologs for agronomically important genes in sweet sorghum. However, the functions of these genes are to be evaluated as most of them are uni-di exonic on SSRG. GWAS studies may help to understand their role further.

## Availability of data and materials

The superTranscriptome assembly is publicly available at the European Nucleotide Archive (ENA) [https://www.ebi.ac.uk/ena/browser/home] with accession number <u>GCA_963506585</u>. Improved genome annotations (*.gff3 files) for 15 diverse sorghum genomes, which include new gene additions and splicing updates using superTranscriptome with PASA gene structure annotation pipeline, will be provided upon request. The R and Python codes used in the present analysis are publicly available on GitHub with repositories named Sorghum superTranscriptome [https://github.com/nikhilshinde0909/Sorghum-superTranscriptome] and Modified necklace pipeline [https://github.com/nikhilshinde0909/Modified-Necklace-Pipeline]. The supplemental data and tables have been submitted along with the manuscript.

## Acknowledgments

We want to thank SRM Institute of Science and Technology, Kattankulathur, Chennai (India), for providing fellowships and the necessary infrastructure for conducting present research work.

## Author contributions

1. Conceptualization: NSR, NS

2. Methodology: NS, SKM

3. Investigation: NSR, NS

4. Visualization: NSR, NS, SKM

5. Supervision: NSR, SKM

6. Writing-original draft: NS

7. Writing-review & editing: NSR, NS

## Abbreviations

PAV: Presence/absence variation
ePAV: Expression presence/absence variation
gPAV: Genomic presence/absence variation
SV: Sources of variation
RTAs: Representative transcript assemblies
CNV: Copy number variation
HISAT2: Hierarchical indexing for spliced alignment of transcripts version 2
BUSCO: Benchmarking universal single-copy orthologs
CPC2: Coding potential calculator version 2
GFF3: General feature format type 3
SAM: Sequence alignment map
BAM: Binary alignment map
LHCs: Light harvesting complexes
DGE: Differential gene expression
DEG: Differentially expressed genes
DEU: Differential exon usage
NCBI: National center for biotechnology information
SRA: Sequence read archives
aa: Amino acid
bp: Base pairs

## References

Agarwala, S. C. et al. (1993) ‘Sugar-cane response to copper in refined sand’, TROP.AGRIC., 70(4).

Ananda, G. K. S. et al. (2020) ‘Wild Sorghum as a Promising Resource for Crop Improvement’, Frontiers in Plant Science. doi: 10.3389/fpls.2020.01108.

Anders, S., Reyes, A. and Huber, W. (2012) ‘Detecting differential usage of exons from RNA-seq data’, Genome Research, 22(10). doi: 10.1101/gr.133744.111.

Aoki, N. et al. (2003) ‘The sucrose transporter gene family in rice’, Plant and Cell Physiology, 44(3). doi: 10.1093/pcp/pcg030.

Babst, B. A. et al. (2021) ‘Physiology and whole-plant carbon partitioning during stem sugar accumulation in sweet dwarf sorghum’, Planta, 254(4). doi: 10.1007/s00425-021-03718-w.

Bhatti, A. et al. (2020) ‘Pan-transcriptomics and its applications’, in Pan-genomics: Applications, Challenges, and Future Prospects. doi: 10.1016/b978-0-12-817076-2.00018-4.

Blighe K, Rana S and Lewis M (2022) Publication-ready volcano plots with enhanced colouring and labeling.

Bodenhofer, U. et al. (2015) ‘Msa: An R package for multiple sequence alignment’, Bioinformatics, 31(24). doi: 10.1093/bioinformatics/btv494.

Boutet, E. et al. (2016) ‘Uniprotkb/swiss-prot, the manually annotated section of the uniprot knowledgebase: How to use the entry view’, in Methods in Molecular Biology. doi: 10.1007/978-1-4939-3167-5_2.

Burks, P. S. et al. (2015) ‘Genomewide association for sugar yield in sweet sorghum’, Crop Science, 55(5). doi: 10.2135/cropsci2015.01.0057.

Chen, S. et al. (2018) ‘Fastp: An ultra-fast all-in-one FASTQ preprocessor’, in Bioinformatics. doi: 10.1093/bioinformatics/bty560.

Contreras, R. A. et al. (2018) ‘Copper stress induces antioxidant responses and accumulation of sugars and phytochelatins in Antarctic Colobanthus quitensis (Kunth) Bartl.’, Biological Research, 51(1). doi: 10.1186/s40659-018-0197-0.

Cooper, E. A. et al. (2019) ‘A new reference genome for Sorghum bicolor reveals high levels of sequence similarity between sweet and grain genotypes: implications for the genetics of sugar metabolism’, BMC Genomics 2019 20:1. BioMed Central, 20(1), pp. 1–doi: 10.1186/S12864-019-5734-X.

Dai, X. et al. (2013) ‘PlantTFcat: An online plant transcription factor and transcriptional regulator categorization and analysis tool’, BMC Bioinformatics, 14(1). doi: 10.1186/1471-2105-14-321.

Davidson, N. M., Hawkins, A. D. K. and Oshlack, A. (2017) ‘SuperTranscripts: A data driven reference for analysis and visualisation of transcriptomes’, Genome Biology, 18(1). doi: 10.1186/s13059-017-1284-1.

Davidson, N. M. and Oshlack, A. (2014) ‘Corset: Enabling differential gene expression analysis for de novo assembled transcriptomes’, Genome Biology, 15(7). doi: 10.1186/s13059-014-0410-6.

Davidson, N. M. and Oshlack, A. (2018) ‘Necklace: combining reference and assembled transcriptomes for more comprehensive RNA-Seq analysis’, GigaScience, 7(5). doi: 10.1093/gigascience/giy045.

Deol, K. K. et al. (2013) ‘Identification and characterization of the three homeologues of a new sucrose transporter in hexaploid wheat (Triticum aestivum L.)’, BMC Plant Biology, doi: 10.1186/1471-2229-13-181.

Ellstrand, N. C., Prentice, H. C. and Hancock, J. F. (1999) ‘Gene flow and introgression from domesticated plants into their wild relatives’, Annual Review of Ecology and Systematics, 30. doi: 10.1146/annurev.ecolsys.30.1.539.

F., C., et al. (2006) ‘Sequencing Medicago truncatula expressed sequenced tags using 454 Life Sciences technology.’, BMC Genomics, 7.

Gerdol, M. et al. (2020) ‘Massive gene presence-absence variation shapes an open pan-genome in the Mediterranean mussel’, Genome biology, 21(1). doi: 10.1186/s13059-020-02180-3.

Ghaffari, N. et al. (2014) ‘Novel transcriptome assembly and improved annotation of the whiteleg shrimp (Litopenaeus vannamei), a dominant crustacean in global seafood mariculture’, Scientific Reports, 4. doi: 10.1038/srep07081.

Gómez-Rubio, V. (2017) ‘ggplot2 - Elegant Graphics for Data Analysis (2nd Edition)’, Journal of Statistical Software, 77(Book Review 2). doi: 10.18637/jss.v077.b02.

Grabherr, M. G. et al. (2011) ‘Full-length transcriptome assembly from RNA-Seq data without a reference genome’, Nature Biotechnology, 29(7). doi: 10.1038/nbt.1883.

Guo, A. Y. et al. (2008) ‘PlantTFDB: A comprehensive plant transcription factor database’, Nucleic Acids Research, 36(SUPPL. 1). doi: 10.1093/nar/gkm841.

Haas, B. J. et al. (2008) ‘Automated eukaryotic gene structure annotation using EVidenceModeler and the Program to Assemble Spliced Alignments’, Genome Biology, 9(1). doi: 10.1186/gb-2008-9-1-r7.

Hao, Z. et al. (2020) ‘RIdeogram: Drawing SVG graphics to visualize and map genome-wide data on the idiograms’, PeerJ Computer Science, 6. doi: 10.7717/peerj-cs.251.

Hirose, T. et al. (2010) ‘Disruption of a gene for rice sucrose transporter, OsSUT1, impairs pollen function but pollen maturation is unaffected’, Journal of Experimental Botany, 61(13). doi: 10.1093/jxb/erq175.

Hirsch, C. N. et al. (2014) ‘Insights into the maize pan-genome and pan-transcriptome’, Plant Cell, 26(1). doi: 10.1105/tpc.113.119982.

Hu, Y. et al. (2022) ‘Genome-wide identification of chromatin regulators in Sorghum bicolor’, 3 Biotech, 12(5). doi: 10.1007/s13205-022-03181-8.

Hufford, M. B. et al. (2013) ‘The Genomic Signature of Crop-Wild Introgression in Maize’, PLoS Genetics, 9(5). doi: 10.1371/journal.pgen.1003477.

Hufford, M. B. et al. (2021) ‘De novo assembly, annotation, and comparative analysis of 26 diverse maize genomes’, Science, 373(6555). doi: 10.1126/science.abg5289.

Jayakodi, M. et al. (2020) ‘The barley pan-genome reveals the hidden legacy of mutation breeding’, Nature, 588(7837). doi: 10.1038/s41586-020-2947-8.

Jiao, W. B. and Schneeberger, K. (2017) ‘The impact of third generation genomic technologies on plant genome assembly’, Current Opinion in Plant Biology. doi: 10.1016/j.pbi.2017.02.002.

Jin, J. et al. (2021) ‘PLncDB V2.0: A comprehensive encyclopedia of plant long noncoding RNAs’, Nucleic Acids Research, 49(D1). doi: 10.1093/nar/gkaa910.

Jin, M. et al. (2016) ‘Maize pan-transcriptome provides novel insights into genome complexity and quantitative trait variation’, Scientific Reports, 6. doi: 10.1038/srep18936.

Jobson, E. and Roberts, R. (2022) ‘Genomic structural variation in tomato and its role in plant immunity’, Molecular Horticulture, 2(1). doi: 10.1186/s43897-022-00029-w.

Kadier, Y. et al. (2017) ‘Genome-wide identification, classification and expression analysis of NAC family of genes in sorghum [Sorghum bicolor (L.) Moench]’, Plant Growth Regulation, 83(2). doi: 10.1007/s10725-017-0295-y.

Kang, Y. J. et al. (2017) ‘CPC2: A fast and accurate coding potential calculator based on sequence intrinsic features’, Nucleic Acids Research, 45(W1). doi: 10.1093/nar/gkx428.

Kent, W. J. (2002) ‘ BLAT —The BLAST-Like Alignment Tool’, Genome Research, 12(4). doi: 10.1101/gr.229202.

Kim, D. et al. (2019) ‘Graph-based genome alignment and genotyping with HISAT2 and HISAT-genotype’, Nature Biotechnology, 37(8). doi: 10.1038/s41587-019-0201-4.

Leach, K. A. et al. (2017) ‘Sucrose transporter2 contributes to maize growth, development, and crop yield’, Journal of Integrative Plant Biology, 59(6). doi: 10.1111/jipb.12527.

Li, C. et al. (2019) ‘The HuangZaoSi Maize Genome Provides Insights into Genomic Variation and Improvement History of Maize’, Molecular Plant, 12(3). doi: 10.1016/j.molp.2019.02.009.

Li, H. et al. (2009) ‘The Sequence Alignment/Map format and SAMtools’, Bioinformatics, 25(16). doi: 10.1093/bioinformatics/btp352.

Li, Xiaoxia et al. (2014) ‘Molecular characterization and expression patterns of sucrose transport-related genes in sweet sorghum under defoliation’, Acta Physiologiae Plantarum, 36(5). doi: 10.1007/s11738-014-1505-0.

Li, Y. et al. (2019) ‘Transcriptome and metabolome reveal distinct carbon allocation patterns during internode sugar accumulation in different sorghum genotypes’, Plant Biotechnology Journal, 17(2). doi: 10.1111/pbi.12991.

Liao, Y., Smyth, G. K. and Shi, W. (2014) ‘FeatureCounts: An efficient general purpose program for assigning sequence reads to genomic features’, Bioinformatics, 30(7). doi: 10.1093/bioinformatics/btt656.

Liu, Z., Fan, H. and Ma, Z. (2022) ‘Comparison of SWEET gene family between maize and foxtail millet through genomic, transcriptomic, and proteomic analyses’, Plant Genome, 15(3), pp. 1–25. doi: 10.1002/tpg2.20226.

Llerena, J. P. P. et al. (2021) ‘Metallothionein production is a common tolerance mechanism in four species growing in polluted Cu mining areas in Peru’, Ecotoxicology and Environmental Safety, 212. doi: 10.1016/j.ecoenv.2021.112009.

Love, M. I., Huber, W. and Anders, S. (2014) ‘Moderated estimation of fold change and dispersion for RNA-seq data with DESeq2’, Genome Biology, 15(12). doi: 10.1186/s13059-014-0550-8.

Lv, Y. et al. (2019) ‘Maize transposable elements contribute to long non-coding RNAs that are regulatory hubs for abiotic stress response’, BMC Genomics, 20(1). doi: 10.1186/s12864-019-6245-5.

Ma, Y. et al. (2019) ‘A pan-transcriptome analysis shows that disease resistance genes have undergone more selection pressure during barley domestication 06 Biological Sciences 0604 Genetics’, BMC Genomics, 20(1). doi: 10.1186/s12864-018-5357-7.

McCormick, R. F. et al. (2018) ‘The Sorghum bicolor reference genome: improved assembly, gene annotations, a transcriptome atlas, and signatures of genome organization’, Plant Journal, 93(2). doi: 10.1111/tpj.13781.

Medina, C. A., Samac, D. A. and Yu, L. X. (2021) ‘Pan-transcriptome identifying master genes and regulation network in response to drought and salt stresses in Alfalfa (Medicago sativa L.)’, Scientific Reports, 11(1). doi: 10.1038/s41598-021-96712-x.

Mizuno, H., Kasuga, S. and Kawahigashi, H. (2016) ‘The sorghum SWEET gene family: Stem sucrose accumulation as revealed through transcriptome profiling’, Biotechnology for Biofuels, 9(1). doi: 10.1186/s13068-016-0546-6.

Moriya, Y. et al. (2007) ‘KAAS: An automatic genome annotation and pathway reconstruction server’, Nucleic Acids Research, 35(SUPPL.2). doi: 10.1093/nar/gkm321.

Pagès, H. et al. (2022) ‘AnnotationDbi: Manipulation of SQLite-based annotations in Bioconductor’, R package version, 1.

Pertea, G. and Pertea, M. (2020) ‘GFF Utilities: GffRead and GffCompare’, F1000Research, 9. doi: 10.12688/f1000research.23297.2.

Pertea, M. et al. (2015) ‘StringTie enables improved reconstruction of a transcriptome from RNA-seq reads’, Nature Biotechnology, 33(3). doi: 10.1038/nbt.3122.

Petek, M. et al. (2020) ‘Cultivar-specific transcriptome and pan-transcriptome reconstruction of tetraploid potato’, Scientific Data, 7(1). doi: 10.1038/s41597-020-00581-4.

Punia, H. et al. (2021) ‘Genome-wide transcriptome profiling, characterization, and functional identification of nac transcription factors in sorghum under salt stress’, Antioxidants, 10(10). doi: 10.3390/antiox10101605.

Qin, P. et al. (2021) ‘Pan-genome analysis of 33 genetically diverse rice accessions reveals hidden genomic variations’, Cell, 184(13). doi: 10.1016/j.cell.2021.04.046.

R. Benatti, M., et al. (2014) ‘Metallothionein deficiency impacts copper accumulation and redistribution in leaves and seeds of Arabidopsis’, New Phytologist, 202(3). doi: 10.1111/nph.12718.

Radchuk, V. et al. (2017) ‘Down-regulation of the sucrose transporters HvSUT1 and HvSUT2 affects sucrose homeostasis along its delivery path in barley grains’, Journal of Experimental Botany, 68(16). doi: 10.1093/jxb/erx266.

Rao, P. S. et al. (2019) ‘Sorghum: A multipurpose bioenergy crop’, in Sorghum: State of the Art and Future Perspectives. doi: 10.2134/agronmonogr58.2014.0074.

Ruperao, P. et al. (2021) ‘Sorghum Pan-Genome Explores the Functional Utility for Genomic-Assisted Breeding to Accelerate the Genetic Gain’, Frontiers in Plant Science, 12. doi: 10.3389/fpls.2021.666342.

Sadedin, S. P., Pope, B. and Oshlack, A. (2012) ‘Bpipe: A tool for running and managing bioinformatics pipelines’, Bioinformatics, 28(11). doi: 10.1093/bioinformatics/bts167.

Sakschewski, B. et al. (2014) ‘Feeding 10 billion people under climate change: How large is the production gap of current agricultural systems?’, Ecological Modelling, 288. doi: 10.1016/j.ecolmodel.2014.05.019.

Sanjari, S. et al. (2019) ‘Systematic analysis of NAC transcription factors’ gene family and identification of post-flowering drought stress responsive members in sorghum’, Plant Cell Reports, 38(3). doi: 10.1007/s00299-019-02371-8.

Simão, F. A. et al. (2015) ‘BUSCO: Assessing genome assembly and annotation completeness with single-copy orthologs’, Bioinformatics, 31(19). doi: 10.1093/bioinformatics/btv351.

Smith-Unna, R. et al. (2016) ‘TransRate: Reference-free quality assessment of de novo transcriptome assemblies’, Genome Research, 26(8). doi: 10.1101/gr.196469.115.

Smith, O. et al. (2019) ‘A domestication history of dynamic adaptation and genomic deterioration in Sorghum’, Nature Plants, 5(4). doi: 10.1038/s41477-019-0397-9.

Soneson, C. et al. (2016) ‘Isoform prefiltering improves performance of count-based methods for analysis of differential transcript usage’, Genome Biology, 17(1). doi: 10.1186/s13059-015-0862-3.

Statello, L. et al. (2021) ‘Gene regulation by long non-coding RNAs and its biological functions’, Nature Reviews Molecular Cell Biology. doi: 10.1038/s41580-020-00315-9.

Sui, N. et al. (2015) ‘Identification and transcriptomic profiling of genes involved in increasing sugar content during salt stress in sweet sorghum leaves’, BMC Genomics, 16(1). doi: 10.1186/s12864-015-1760-5.

Sun, C. et al. (2017) ‘RPAN: Rice pan-genome browser for ∼3000 rice genomes’, Nucleic Acids Research, 45(2). doi: 10.1093/nar/gkw958.

Sun, X. et al. (2020) ‘Comparative Transcriptome Analysis Reveals New lncRNAs Responding to Salt Stress in Sweet Sorghum’, Frontiers in Bioengineering and Biotechnology, 8. doi: 10.3389/fbioe.2020.00331.

Sweeney, B. A. et al. (2020) ‘Exploring Non-Coding RNAs in RNAcentral’, Current Protocols in Bioinformatics, 71(1). doi: 10.1002/cpbi.104.

Tao, Y. et al. (2021) ‘Extensive variation within the pan-genome of cultivated and wild sorghum’, Nature Plants, 7(6). doi: 10.1038/s41477-021-00925-x.

Unver, T. and Tombuloglu, H. (2020) ‘Barley long non-coding RNAs (lncRNA) responsive to excess boron’, Genomics, 112(2). doi: 10.1016/j.ygeno.2019.11.007.

Venkateswaran, K., Elangovan, M. and Sivaraj, N. (2018) ‘Origin, domestication and diffusion of Sorghum bicolor’, in Breeding Sorghum for Diverse End Uses. doi: 10.1016/B978-0-08-101879-8.00002-4.

Voelker, W. G. et al. (2023) ‘Ten new high-quality genome assemblies for diverse bioenergy sorghum genotypes’, Frontiers in Plant Science, 13(January), pp. 1–13. doi: 10.3389/fpls.2022.1040909.

Wang, B. et al. (2021) ‘Pan-genome analysis in sorghum highlights the extent of genomic variation and sugarcane aphid resistance genes’, bioRxiv. doi: 10.1101/2021.01.03.424980.

Wang, H. et al. (2015) ‘Analysis of non-coding transcriptome in rice and maize uncovers roles of conserved lncRNAs associated with agriculture traits’, Plant Journal, 84(2). doi: 10.1111/tpj.13018.

Wu, T. et al. (2021) ‘clusterProfiler 4.0: A universal enrichment tool for interpreting omics data’, The Innovation, 2(3). doi: 10.1016/j.xinn.2021.100141.

Xia, J. et al. (2018) ‘A sorghum NAC gene is associated with variation in biomass properties and yield potential’, Plant Direct, 2(7). doi: 10.1002/pld3.70.

Xin, Z. et al. (2021) ‘Sorghum genetic, genomic, and breeding resources’, Planta. doi: 10.1007/s00425-021-03742-w.

Yang, N. et al. (2017) ‘Contributions of Zea mays subspecies mexicana haplotypes to modern maize’, Nature Communications, 8(1). doi: 10.1038/s41467-017-02063-5.

Yang, T. et al. (2022) ‘Improved pea reference genome and pan-genome highlight genomic features and evolutionary characteristics’, Nature Genetics. Springer US, 54(October). doi: 10.1038/s41588-022-01172-2.

Yao, C. et al. (2017) ‘A database for orphan genes in poaceae’, Experimental and Therapeutic Medicine, 14(4). doi: 10.3892/etm.2017.4918.

Yao, W. et al. (2015) ‘Exploring the rice dispensable genome using a metagenome-like assembly strategy’, Genome Biology, 16(1). doi: 10.1186/s13059-015-0757-3.

Zhang, H. et al. (2013) ‘Genome-wide survey and characterization of greenbug induced nac transcription factors in sorghum [Sorghum bicolor (L.) Moench]’, Plant & Animal Genome.

Zhang, L. M. et al. (2018) ‘Sweet sorghum originated through selection of dry, a plant-specific nac transcription factor gene[open]’, Plant Cell, 30(10). doi: 10.1105/tpc.18.00313.

Zhao, Q. et al. (2018) ‘Pan-genome analysis highlights the extent of genomic variation in cultivated and wild rice’, Nature Genetics, 50(2). doi: 10.1038/s41588-018-0041-z.

Zheng, L. Y. et al. (2011) ‘Genome-wide patterns of genetic variation in sweet and grain sorghum (Sorghum bicolor)’, Genome Biology, 12(11). doi: 10.1186/gb-2011-12-11-r114.

Zheng, Y. et al. (2016) ‘iTAK: A Program for Genome-wide Prediction and Classification of Plant Transcription Factors, Transcriptional Regulators, and Protein Kinases’, Molecular Plant. doi: 10.1016/j.molp.2016.09.014.

Zhou, R. et al. (2021) ‘Analysis of Rice Transcriptome Reveals the LncRNA/CircRNA Regulation in Tissue Development’, Rice, 14(1). doi: 10.1186/s12284-021-00455-2.

Zhou, W. et al. (2022) ‘Comparative transcriptome analysis in three sorghum (Sorghum bicolor) cultivars reveal genomic basis of differential seed quality’, Plant Biosystems, 156(1). doi: 10.1080/11263504.2020.1851790.

