## Supporting Information: Appendix S1 for "Unlocking the Genetic Landscape: Enhanced Insights into Sweet Sorghum Genomes through Comprehensive superTranscriptomic Analysis"

### Supporting Information: Appendix S1
### Supplemental Figures

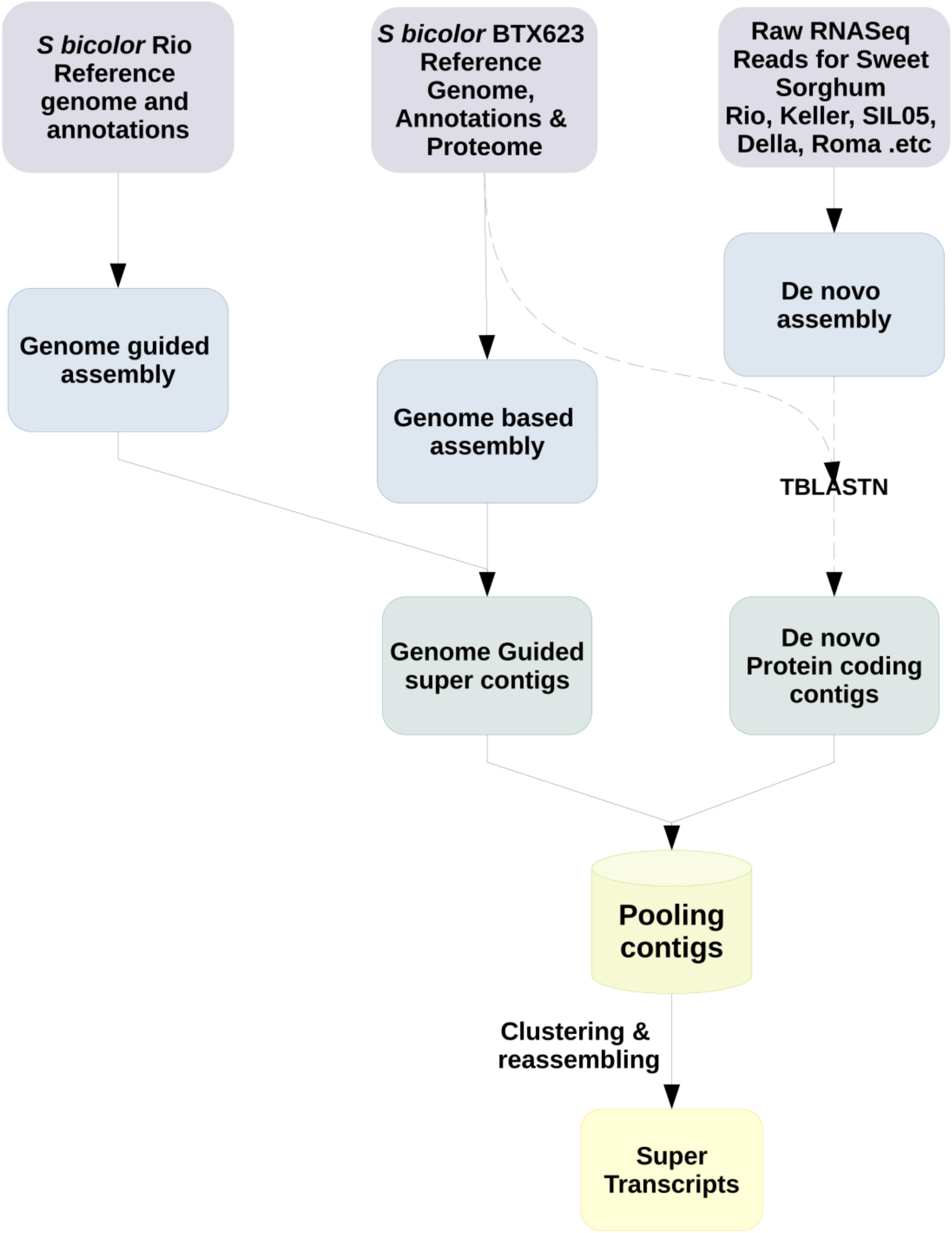


**Supplemental Figure S1**: Necklace pipeline (Davidson and Oshlack, 2018)workflow for 223 RNAseq datasets; combines reference guided and de novo transcriptome assembly approach to get final superTranscripts.

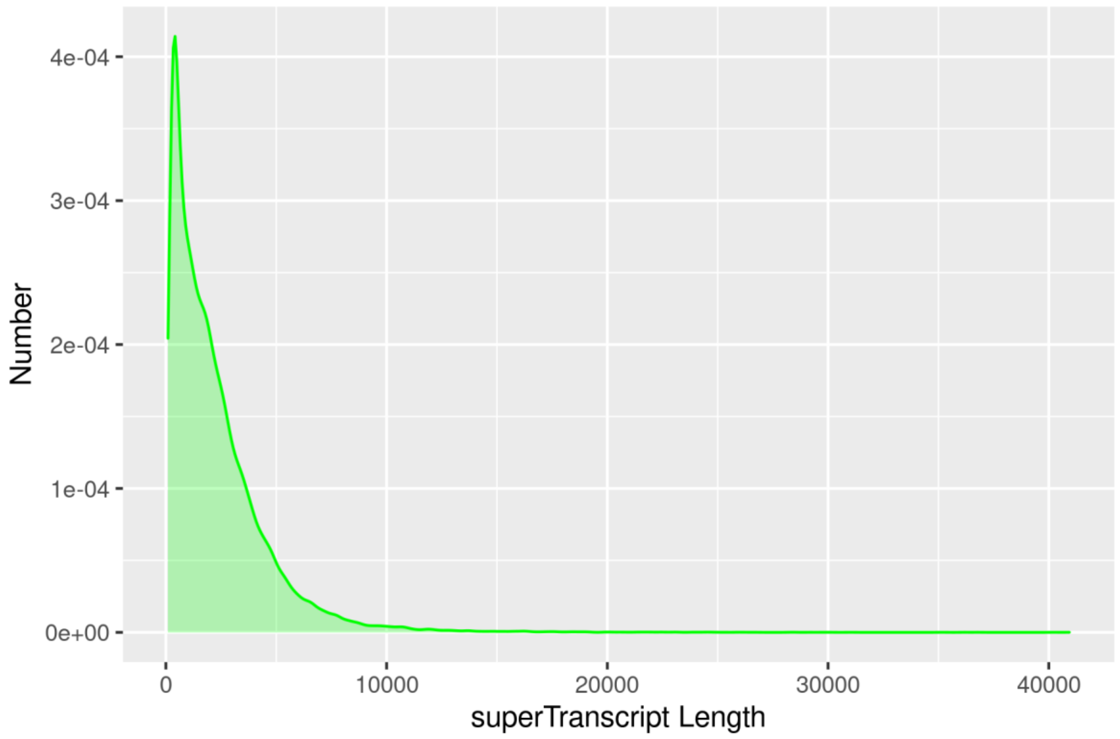


**Supplemental Figure S2**: Density plot showing sweet sorghum superTranscriptome contig length distribution when analyzed with TransRate tool.

| | superTranscriptome | Rio | BTX623 | BTX642 | RTX430 | SC187 | Pangenome |
|---|---|---|---|---|---|---|---|
| Rio-V-M3 | 78.49 | 83.06 | 81.75 | 81.27 | 81.41 | 81.35 | 82.25 |
| Rio-V-M2 | 84.13 | 88.93 | 87.65 | 87.18 | 87.28 | 87.28 | 88.19 |
| Rio-V-M1 | 81.98 | 86.42 | 85.23 | 84.76 | 84.86 | 84.93 | 85.82 |
| Rio-V-L3 | 68.6 | 72.09 | 69.95 | 70.41 | 69.59 | 70.47 | 71.53 |
| Rio-V-I3 | 79.17 | 83.55 | 81.44 | 81.2 | 80.81 | 81.34 | 82.31 |
| Rio-V-I2 | 70.54 | 73.93 | 73.24 | 73.41 | 72.71 | 73.41 | 74.24 |
| Rio-V-I1 | 74.54 | 78.16 | 76.26 | 76.47 | 75.83 | 76.65 | 77.44 |
| Rio-SD-S3 | 77.65 | 82.05 | 79.88 | 80.05 | 79.59 | 79.89 | 80.77 |
| Rio-SD-S2 | 82.52 | 85.04 | 83.55 | 83.67 | 83.33 | 83.76 | 84.62 |
| Rio-SD-S1 | 82.38 | 85.77 | 83.87 | 83.53 | 83.48 | 83.42 | 84.81 |
| Rio-SD-L3 | 73.43 | 78.41 | 76.28 | 76.5 | 75.85 | 76.4 | 77.53 |
| Rio-SD-L2 | 78.47 | 84.64 | 82.52 | 82.23 | 82.07 | 81.97 | 83.6 |
| Rio-SD-L1 | 70.76 | 75.2 | 72.95 | 73.7 | 72.65 | 73.7 | 74.64 |
| Rio-SD-I3 | 86.35 | 92.69 | 91.07 | 90.5 | 90.79 | 90.54 | 91.67 |
| Rio-SD-I2 | 84.43 | 91.05 | 89.12 | 88.56 | 88.76 | 88.49 | 89.81 |
| Rio-SD-I1 | 85.99 | 91.98 | 90.27 | 89.78 | 89.98 | 89.74 | 90.88 |
| Rio-RI-M3 | 81.47 | 85.86 | 84.41 | 83.92 | 84.1 | 84.11 | 85.05 |
| Rio-RI-M2 | 88.76 | 93.49 | 91.9 | 91.53 | 91.58 | 91.71 | 92.61 |
| Rio-RI-M1 | 75.94 | 80.1 | 78.7 | 78.31 | 78.34 | 78.5 | 79.36 |
| Rio-RI-L3 | 79.48 | 83.59 | 82.07 | 81.91 | 81.58 | 81.79 | 82.83 |
| Rio-RI-L2 | 76 | 80.16 | 78.51 | 78.5 | 77.99 | 78.3 | 79.43 |
| Rio-RI-L1 | 83.84 | 88.69 | 87 | 87.19 | 86.69 | 87.12 | 88.17 |
| Rio-RI-I3 | 83.1 | 87.82 | 85.94 | 85.18 | 85.3 | 85.41 | 86.56 |
| Rio-RI-I2 | 74.18 | 78.21 | 76.68 | 76.21 | 76.17 | 76.24 | 77.31 |
| Rio-RI-I1 | 85.99 | 90.59 | 89.08 | 88.65 | 88.66 | 88.7 | 89.71 |
| Rio-FL-M3 | 78.74 | 83.09 | 81.66 | 81.15 | 81.28 | 81.18 | 82.25 |
| Rio-FL-M2 | 80.62 | 85.21 | 83.63 | 83.2 | 83.22 | 83.27 | 84.33 |
| Rio-FL-M1 | 81.31 | 86.38 | 84.62 | 84.11 | 84.16 | 84.18 | 85.39 |
| Rio-FL-L3 | 80.73 | 85.91 | 83.78 | 83.46 | 83.05 | 83.48 | 84.75 |
| Rio-FL-L2 | 85 | 90.35 | 87.32 | 88.99 | 86.85 | 89.17 | 90.26 |
| Rio-FL-L1 | 72.98 | 77.29 | 74.96 | 75.95 | 74.65 | 75.99 | 77.08 |
| Rio-FL-I3 | 84.3 | 89.58 | 87.41 | 86.99 | 86.82 | 86.97 | 88.17 |
| Rio-FL-I2 | 73.24 | 78.2 | 76.04 | 75.73 | 75.33 | 75.7 | 76.85 |
| Rio-FL-I1 | 80.16 | 85.7 | 83.49 | 83 | 83.01 | 83.04 | 84.23 |
| Rio-BL-S2 | 82.55 | 89.24 | 87.8 | 86.99 | 87.42 | 87.1 | 88.64 |
| Rio-BL-S1 | 82.02 | 86.97 | 85.69 | 84.74 | 85.18 | 84.81 | 86.51 |
| Rio-BL-L3 | 78.36 | 86.87 | 84.55 | 84.21 | 84.13 | 84.05 | 85.51 |
| Rio-BL-L2 | 78.48 | 84.09 | 82.22 | 82.05 | 81.89 | 81.92 | 83.13 |
| Rio-BL-L1 | 75.73 | 80.74 | 79.03 | 78.86 | 78.59 | 78.75 | 79.9 |
| Rio-BL-I2 | 85.75 | 93.1 | 91.25 | 90.75 | 90.89 | 90.59 | 91.97 |
| Rio-BL-I1 | 81.7 | 87.25 | 85.56 | 85.07 | 85.26 | 84.98 | 86.15 |
| Rio-ANT-L2 | 77.85 | 83.66 | 81 | 80.89 | 80.46 | 80.56 | 82.1 |
| Rio-ANT-I3 | 87.41 | 93.31 | 91.73 | 91.33 | 91.26 | 91.31 | 92.37 |
| Rio-ANT-I2 | 58.71 | 61.9 | 60.77 | 60.44 | 60.28 | 60.36 | 61.2 |
| Rio-ANT-I1 | 84.72 | 89.79 | 88.01 | 87.62 | 87.4 | 87.54 | 88.67 |
| Rio-ANT-F3 | 85.36 | 91.14 | 89.42 | 89.09 | 89.05 | 88.94 | 90.15 |
| Rio-ANT-F2 | 72.62 | 76.3 | 74.81 | 74.57 | 74.58 | 74.44 | 75.57 |
| Rio-ANT-F1 | 64.51 | 67.73 | 66.1 | 65.84 | 65.74 | 65.8 | 67.02 |

% Read Coverage: 90, 80, 70, 60

**Supplemental Figure S3:** Quality control analysis by aligning raw RNA-seq reads to superTranscriptome and six published sorghum genomes

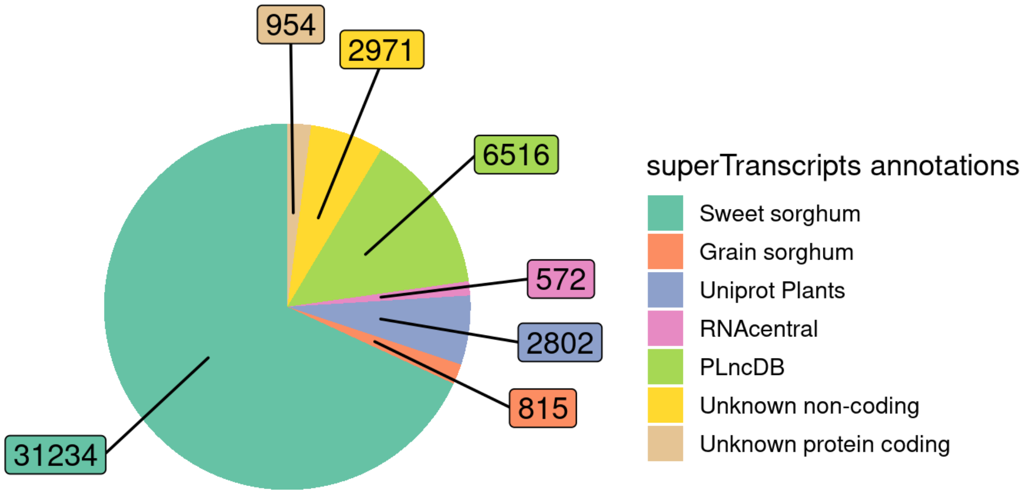


**Supplemental Figure S4:** Pie chart showing showing functional annotations of protein-coding and non-coding sequences in superTranscriptome with different databases.

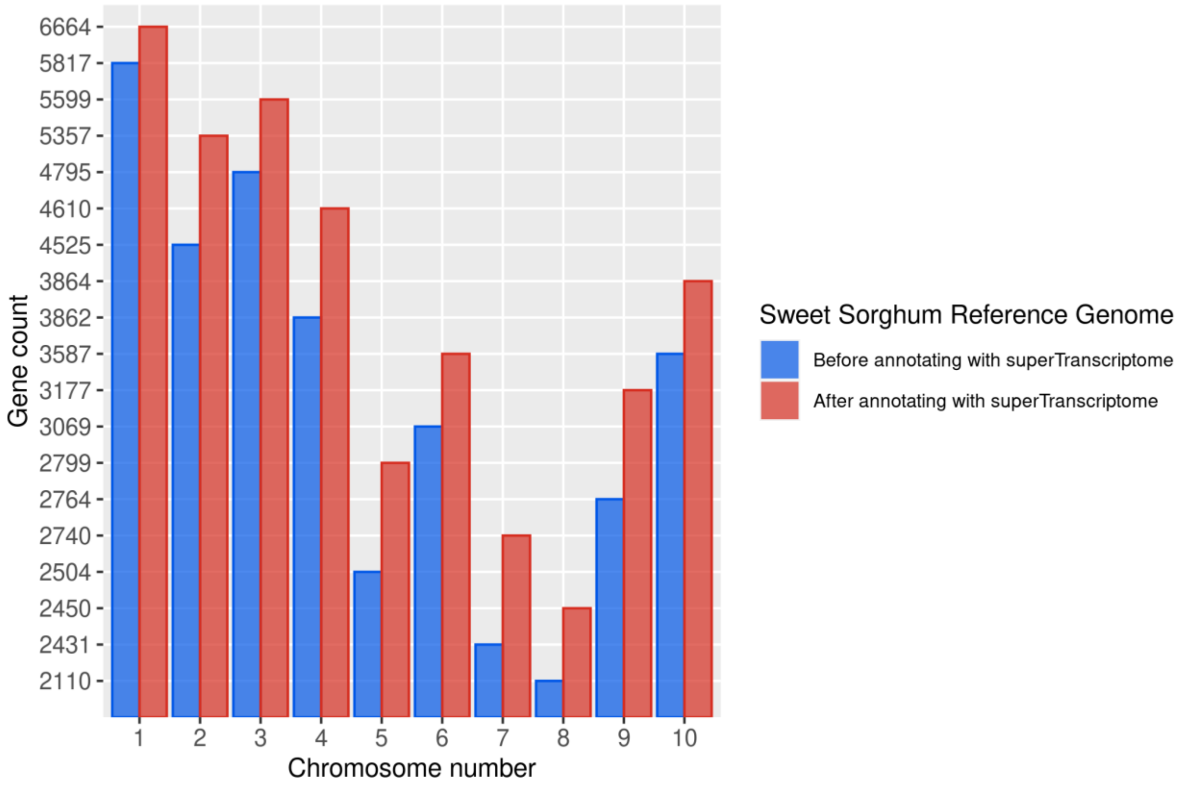


**Supplemental Figure S5:** Chromosome wise number annotated genes on SSRG with superTranscritome by using PASA gene structure annotation pipeline, showing more number of genes are annotated with superTranscriptome based method.

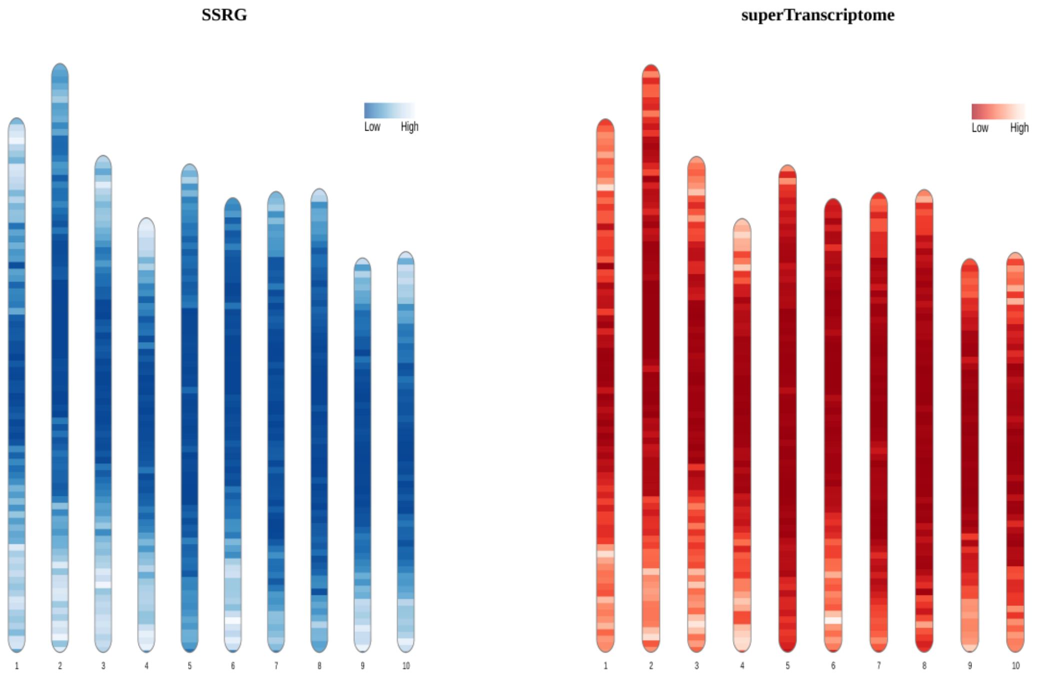


**Supplemental Figure S6: :** Ideogram showing gene densities comparison between SSRG and SSRG annotated with superTranscriptome

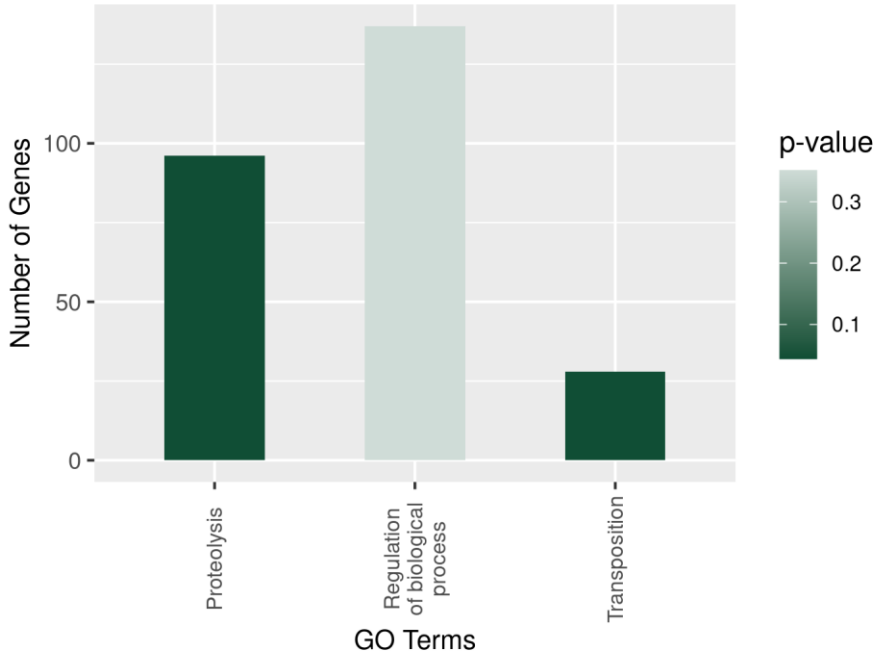


**Supplemental Figure S7(a):** GO enrichment analysis of dispensable genes; suggests that most of the dispensable genes are associated with TEs, proteolysis, and regulation.

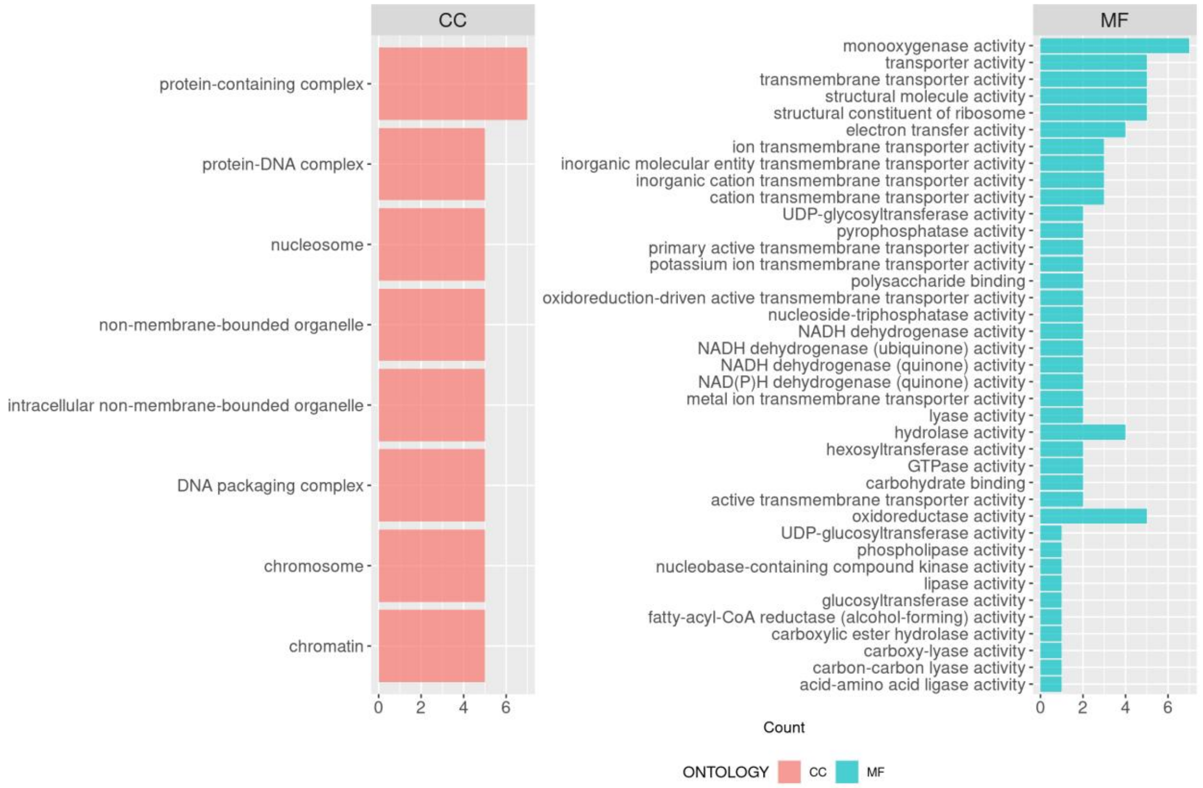


**Supplemental Figure S7(b):** GO terms for 301 genes which are exclusively reported on sweet sorghum Rio, Wray and Leoti cultivar genomes.

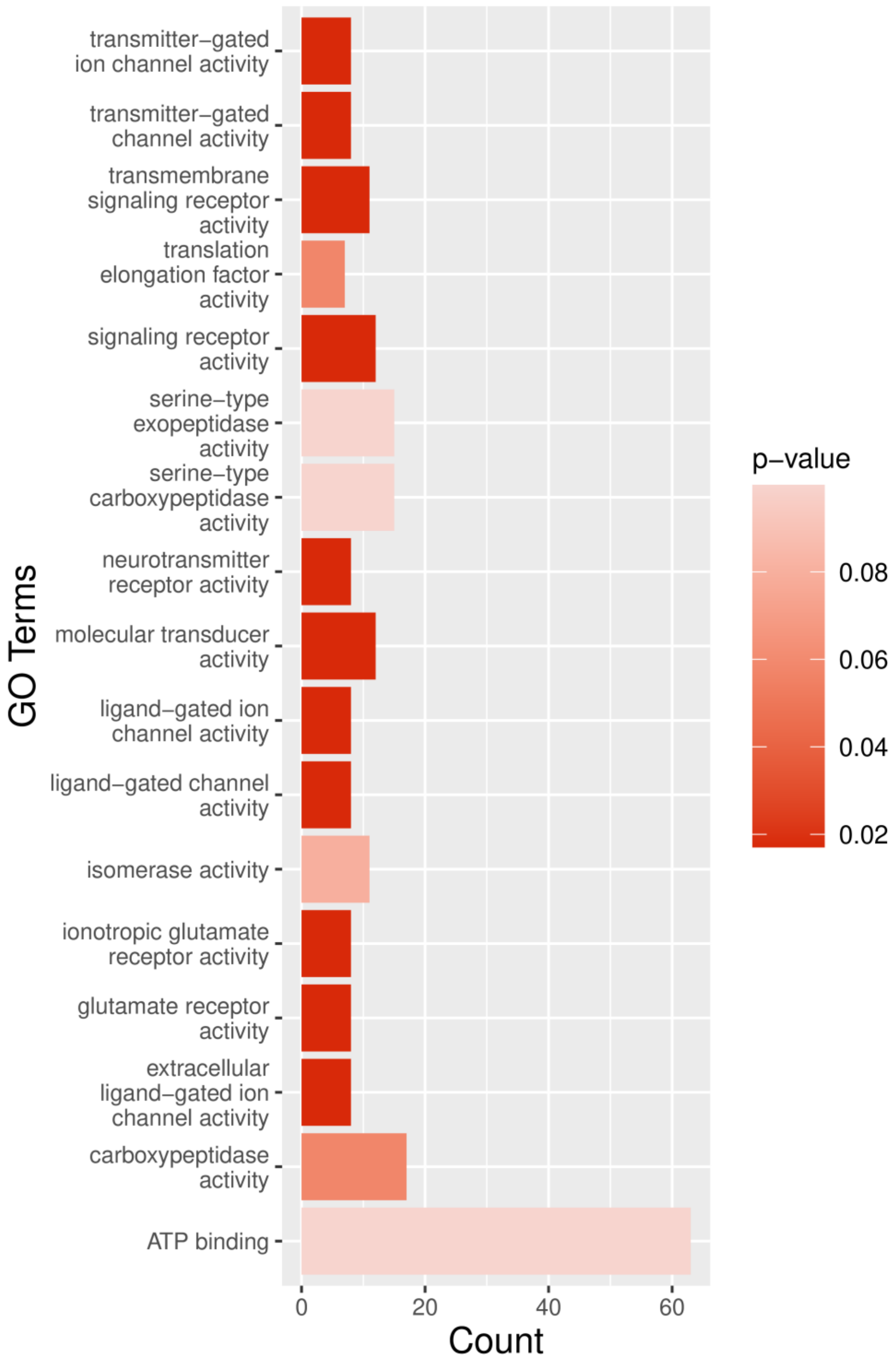


**Supplemental Figure S8(a):** GO enrichment analysis for orphan genes; suggesting that genes associated with cellular signaling are energy metabolism enriched; hence conferring some additional traits too sweet sorghum.

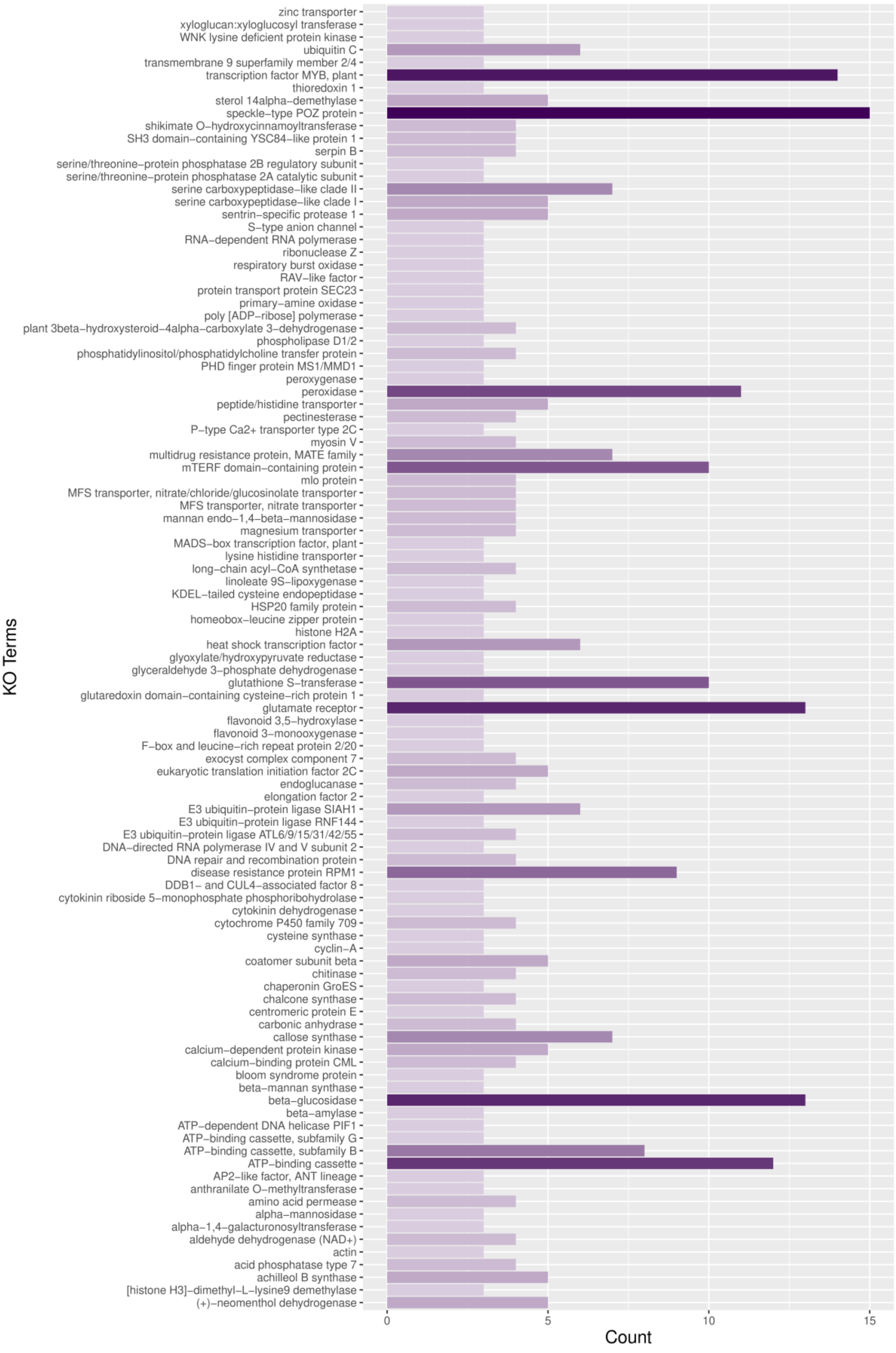


**Supplemental Figure S8(b):** KEGG Orthology (KO) analysis with KAAS reported orphan genes are associated with disease resistance abiotic stress tolerance and cell wall development.

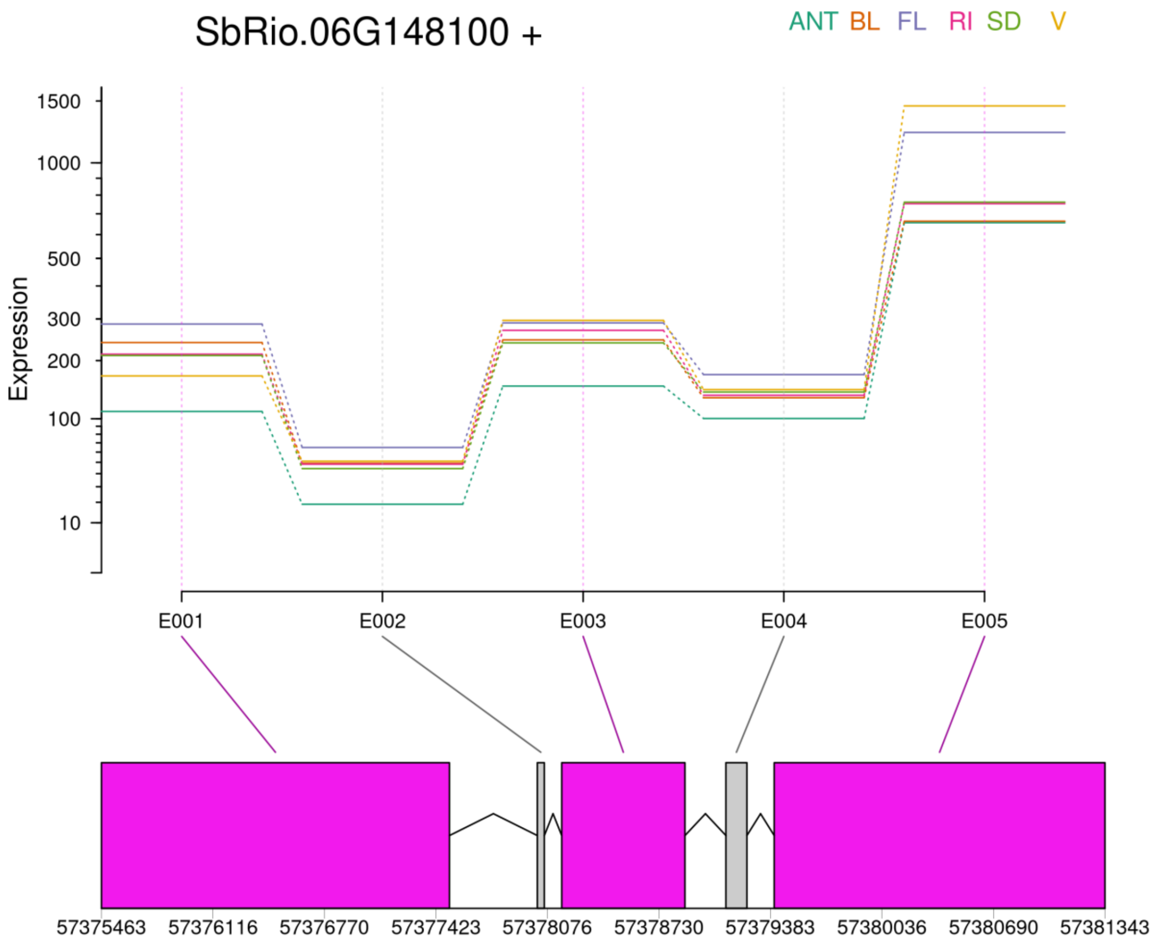


**Supplemental Figure S9(a):** Standard blocking scheme employed for DEU analysis with SSRG yielded 5 exons for the NLP2 TF coding gene, located on chromosome 6 (SbRio.06G148100) of which 3 are differentially used for spliced transcript formation under six developmental stages

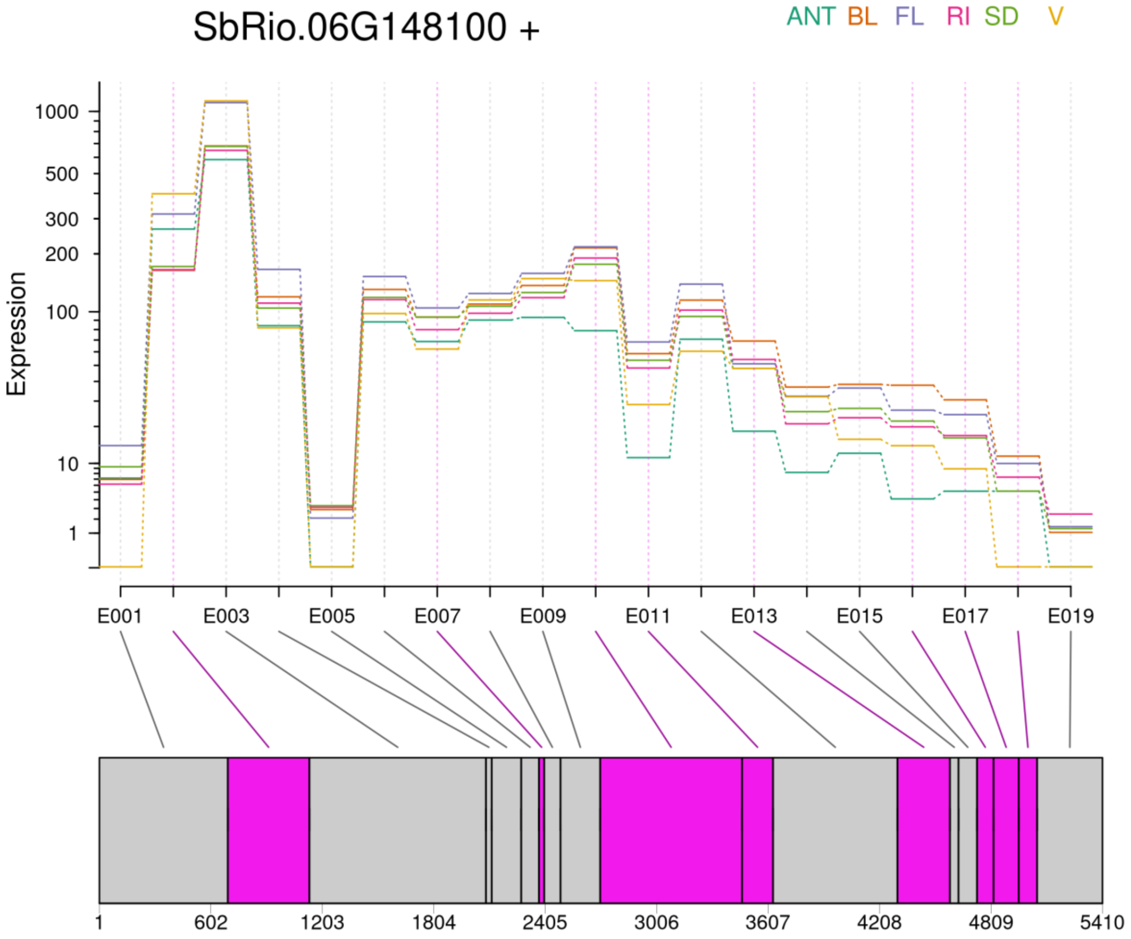


**Supplemental Figure S9(b):** superTranscriptome followed dynamic blocking scheme for same gene (NLP2 TF, gene id: SbRio.06G148100) and identified 19 different exon bins of which 8 are differentially used for alternatively spliced transcript formation under six developmental stages

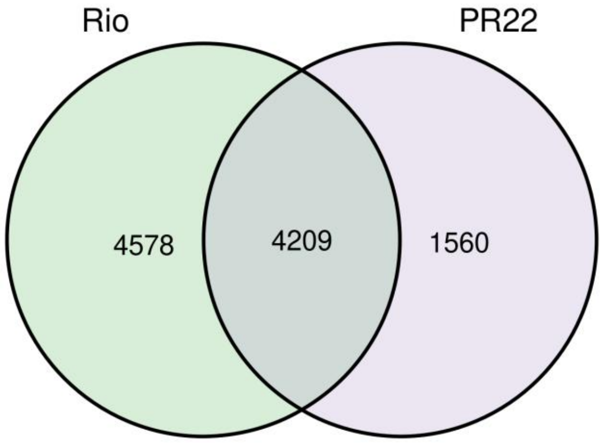


**Supplemental Figure S10:** DEU comparison between Rio and PR22 reported different set of genes that has been differentially used by these two genotypes during internode development; suggesting that differential slicing is extensive in Rio than PR22.

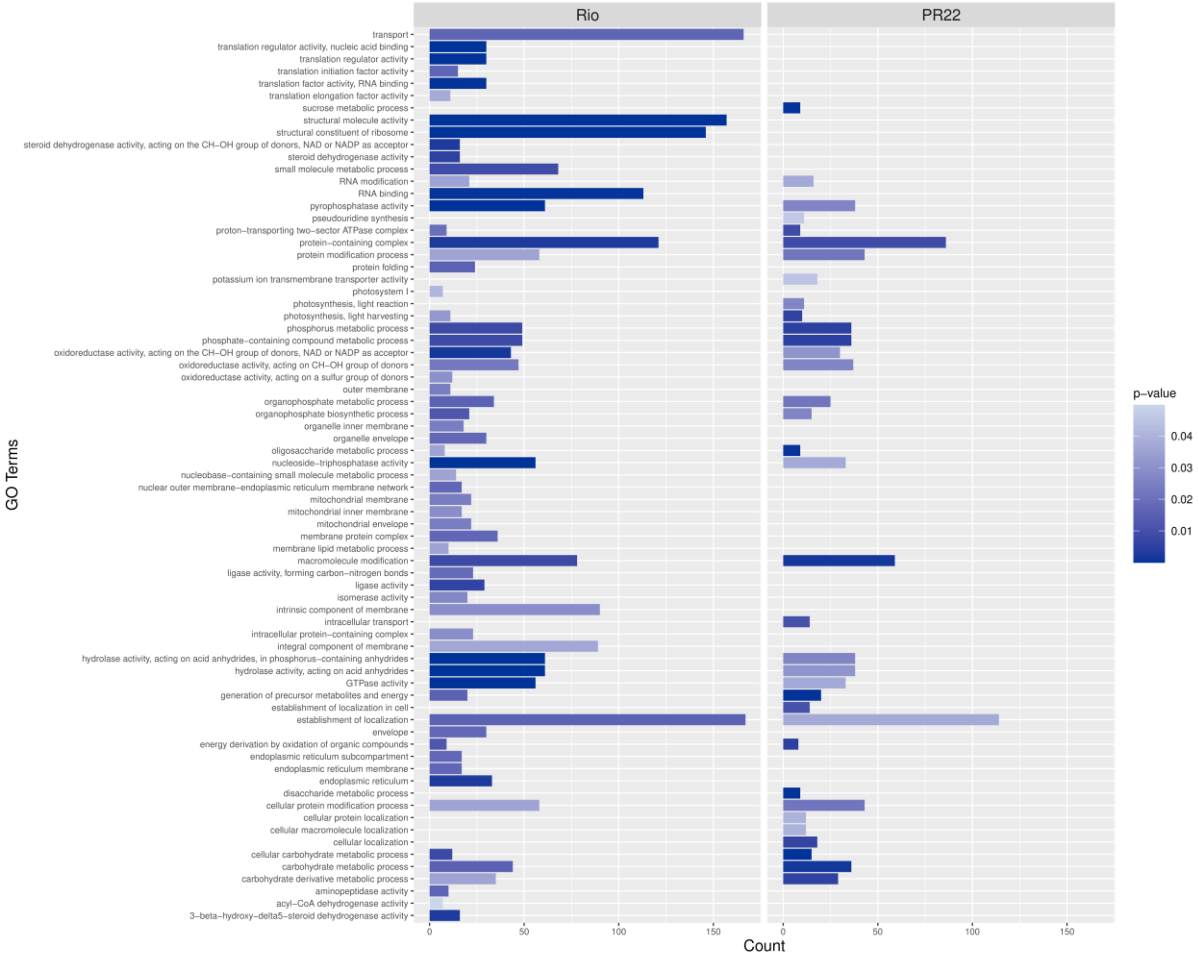


**Supplemental Figure S11:** GO enrichment analysis for differentially spliced genes between Rio and PR22 during internode development

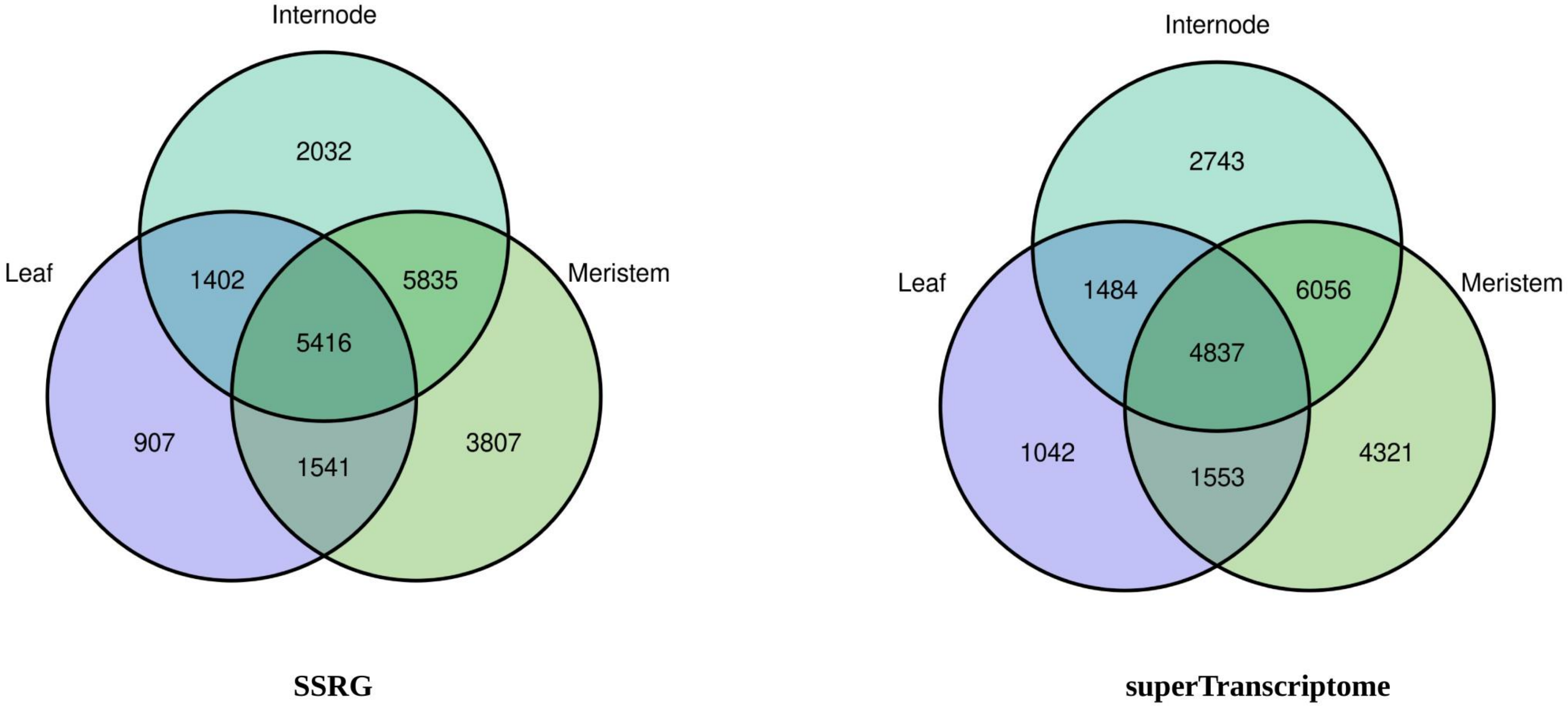


**Supplemental Figure S12(a):** Venn diagram showing number of differentially expressed (P-value < 0.05) genes across Leaf, Internode and Meristem tissues during six time points with two different references.

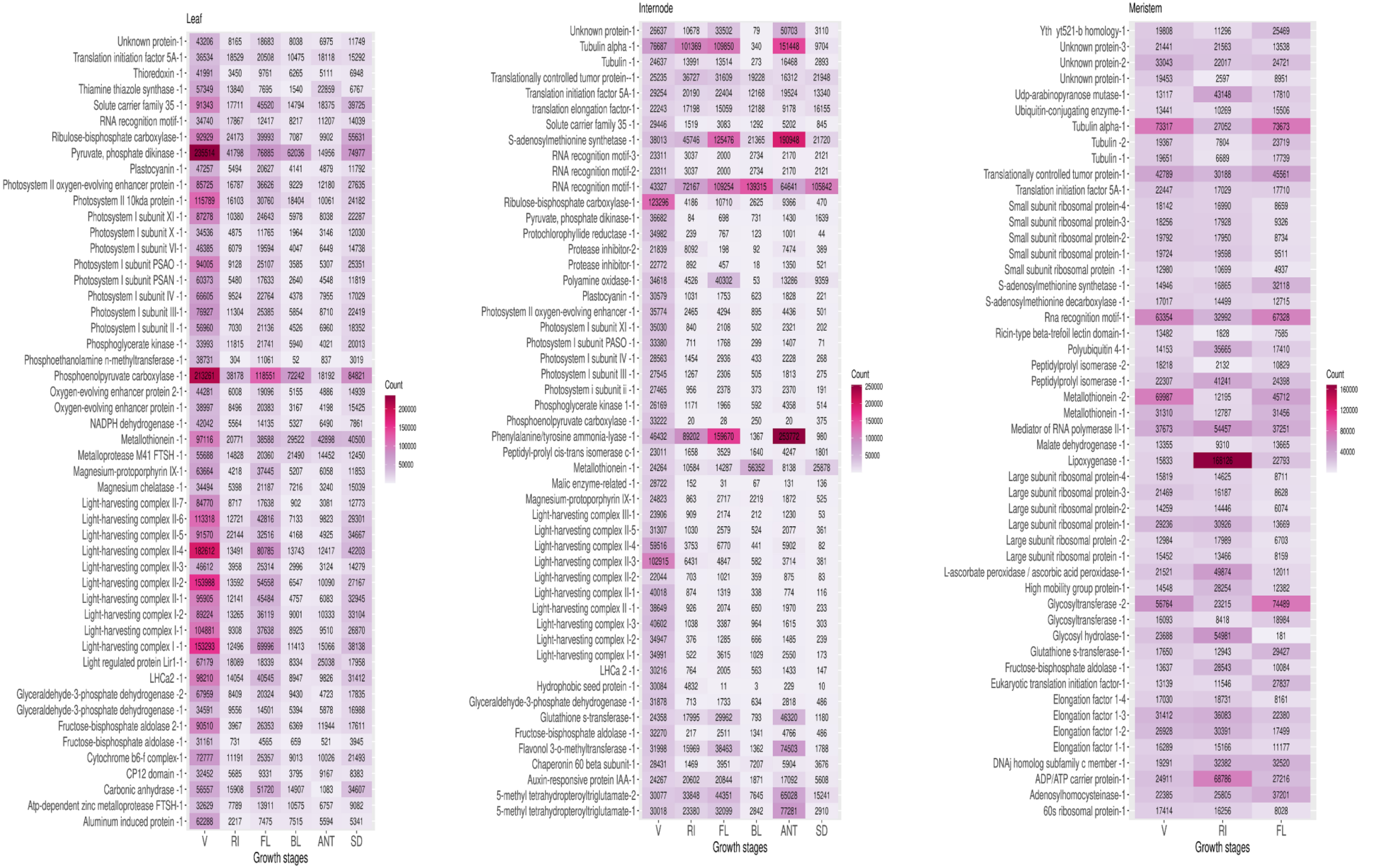


**Supplemental Figure S12(b)**: Heatmap showing top 50 highly expressed genes when SSRG used as a reference for DGE analysis

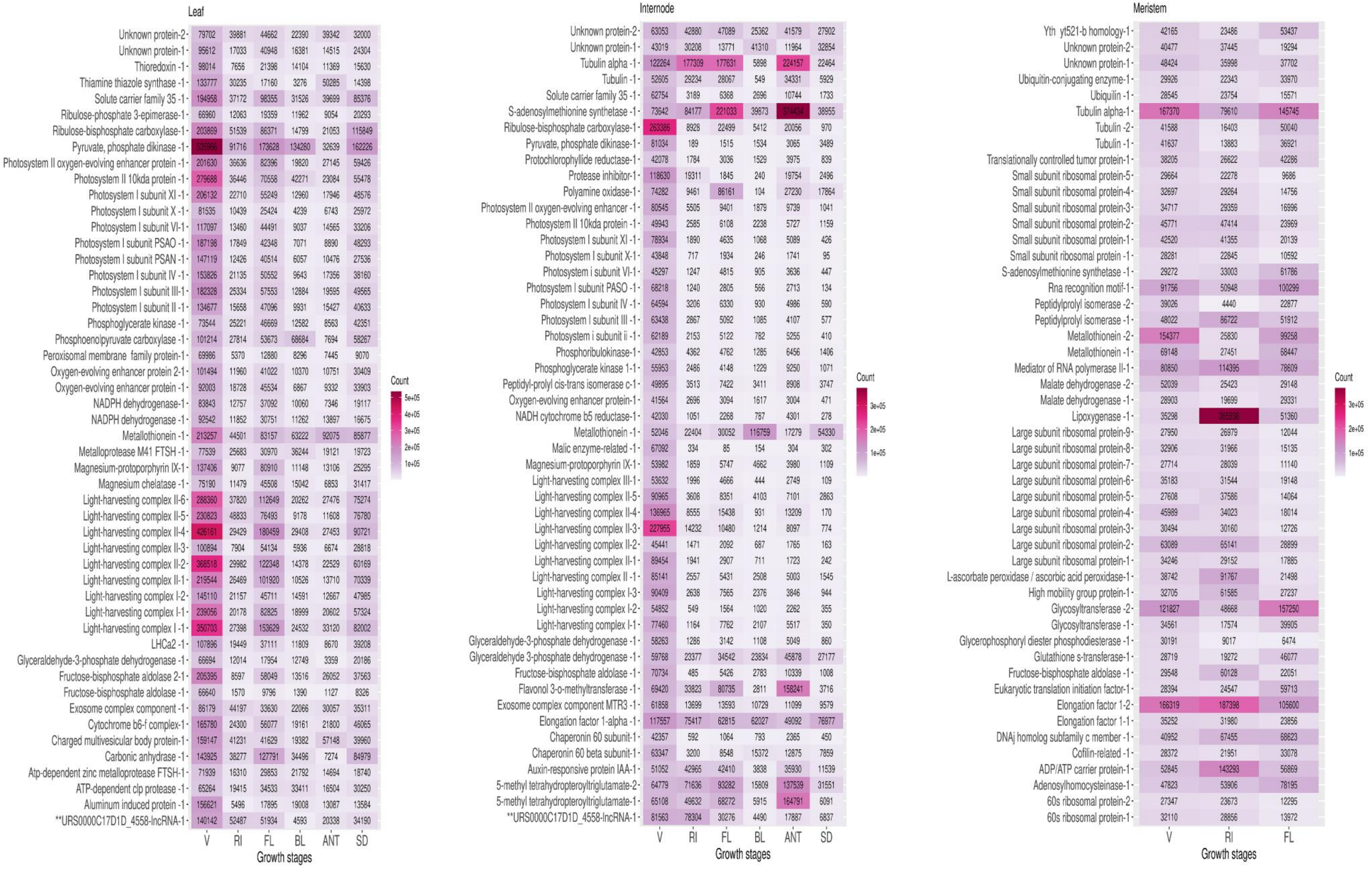


**Supplemental Figure S12(c):** Heatmap showing top 50 highly expressed genes when superTranscriptome used as a reference for DGE analysis

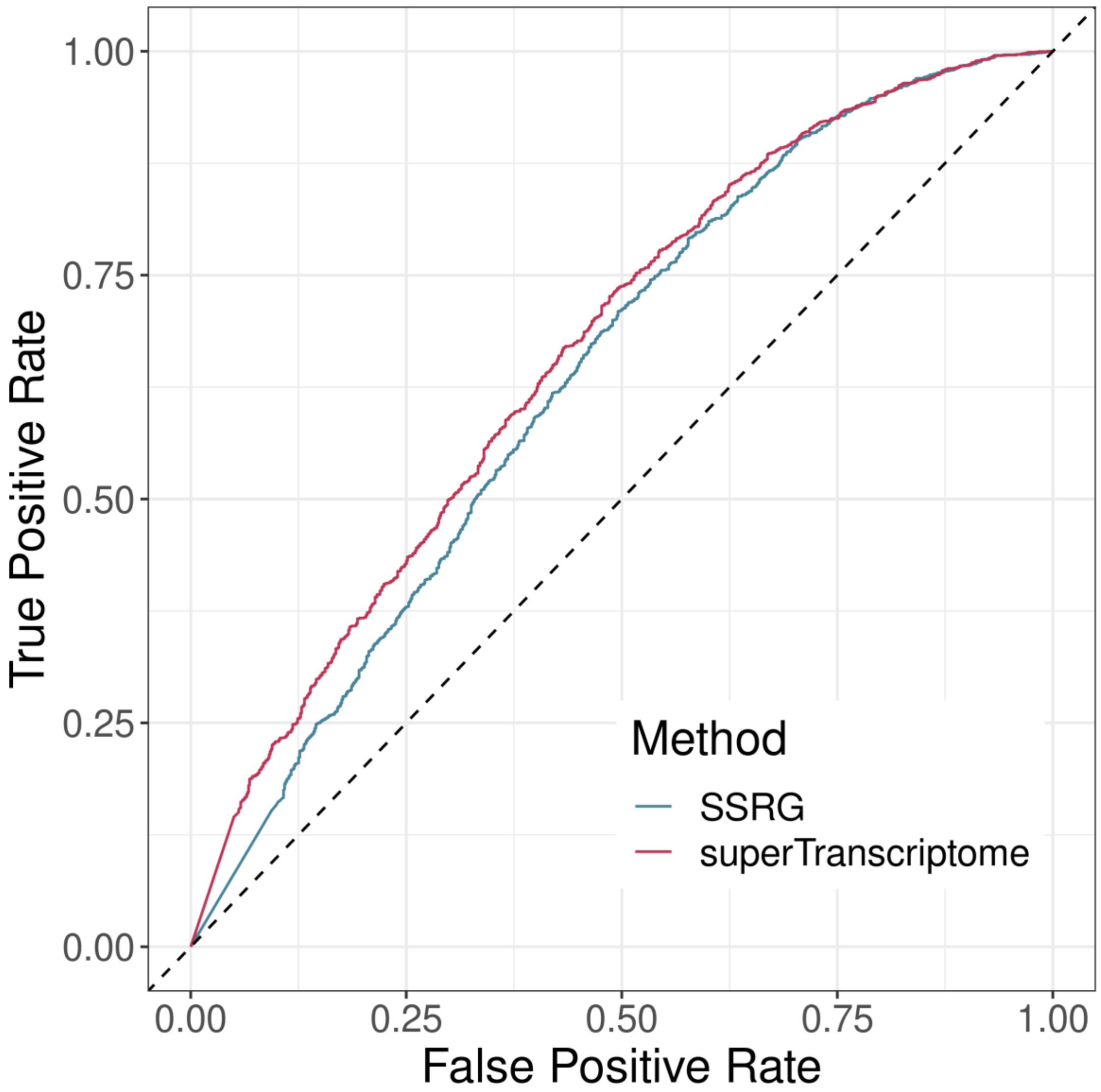


**Supplemental Figure S13:** ROC curve based on TPR and FPR values showing superTranscritptome giving better estimates of DGE than SSRG based standard approach when datasets trained with logistic regression method.

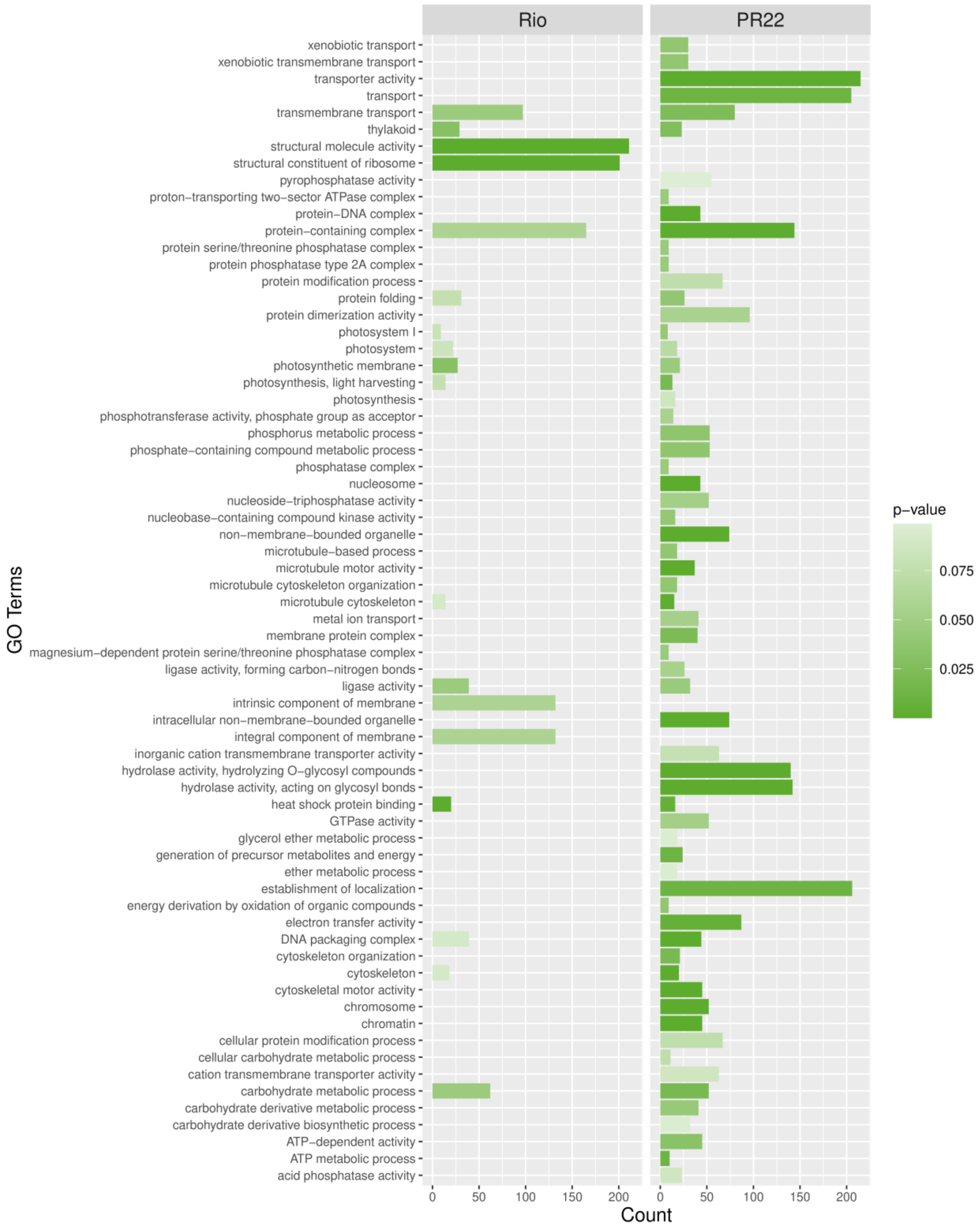


**Supplemental Figure S14:** GO enrichment analysis for DEGs in Rio and PR22 during internode development; reported significant enrichment in GO terms related metal ion transport, hydrolase activity, various transports and secondary metabolite synthesis in PR22

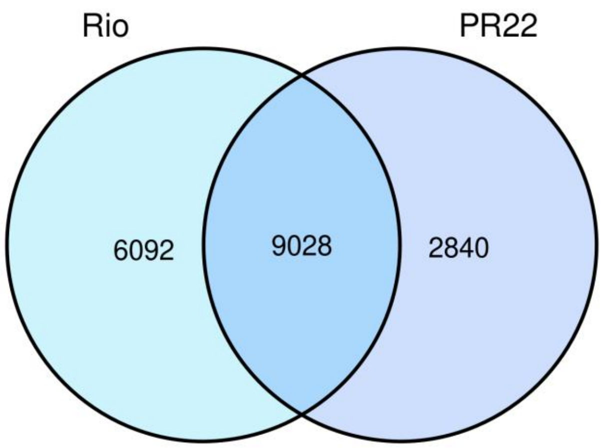


**Supplemental Figure S15:** Venn diagram showing number of differentially expressed genes (P-value < 0.05) in Rio and PR22 during internode development; reported 6092 and 2840 genes are unique to Rio and PR22 respectively.

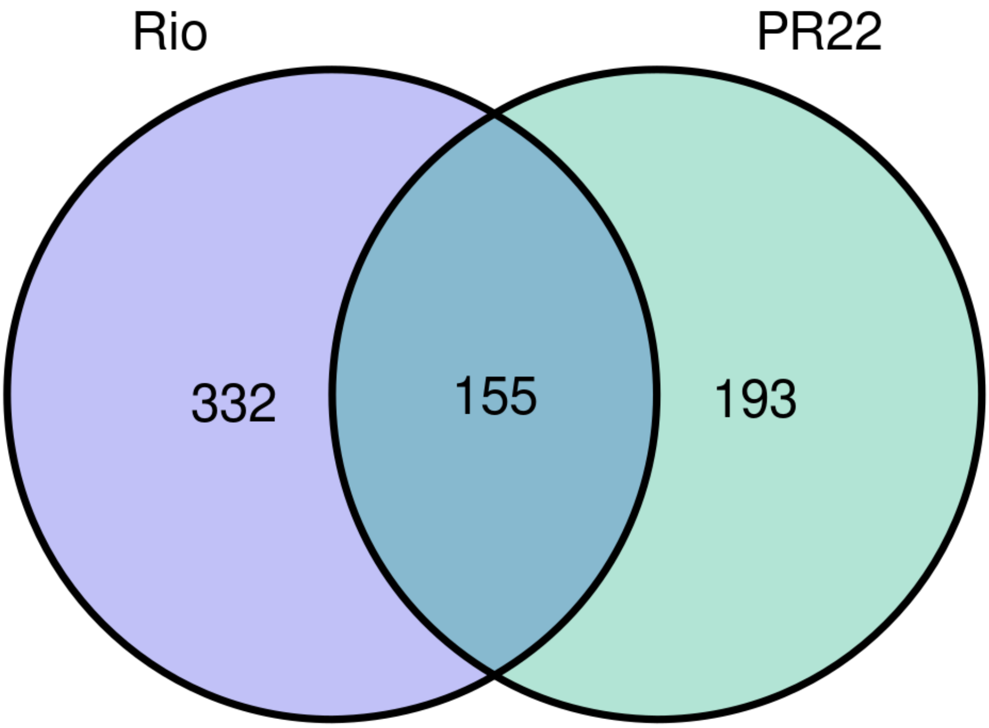


**Supplemental Figure S16(a):** Venn diagram showing number of differentially expressed lncRNAs (P-value < 0.05) in Rio and PR22 during internode development; reported 332 and 193 lncRNAs are unique to Rio and PR22 respectively

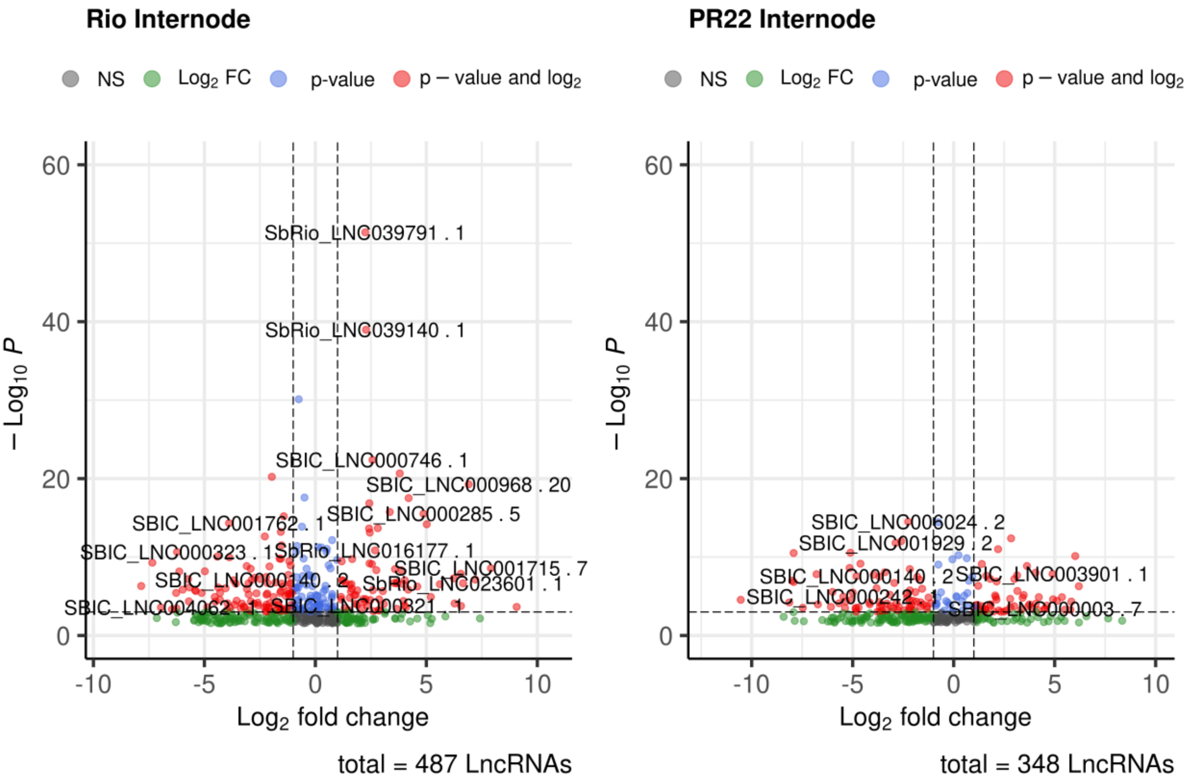


**Supplemental Figure S16(b):** Volcano plot showing lncRNAs reported with contrasting changes in expression during internode development of Rio and PR22 respectively.

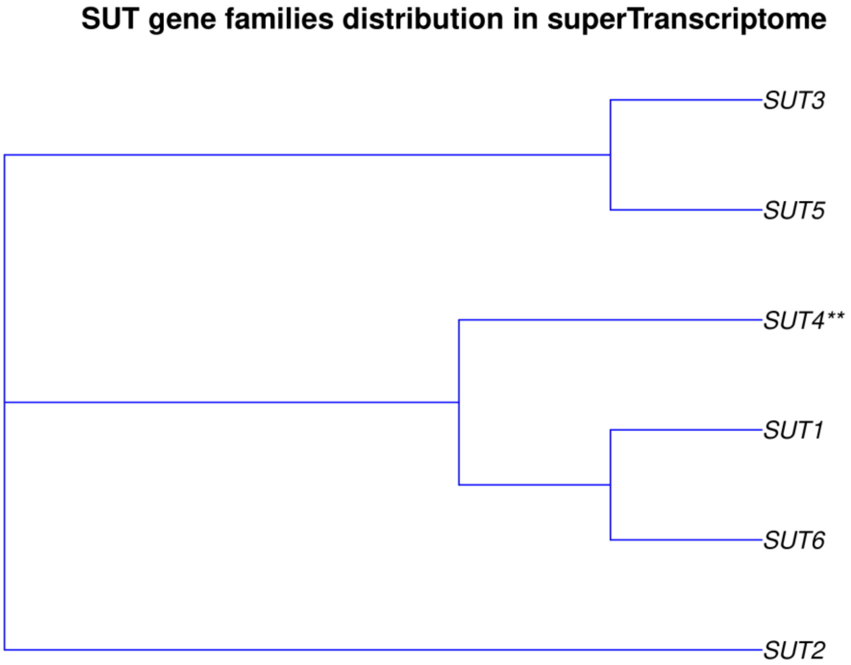


**Supplemental Figure S17:** Multiple Sequence Alignment showing SUT gene families distribution in superTranscriptome

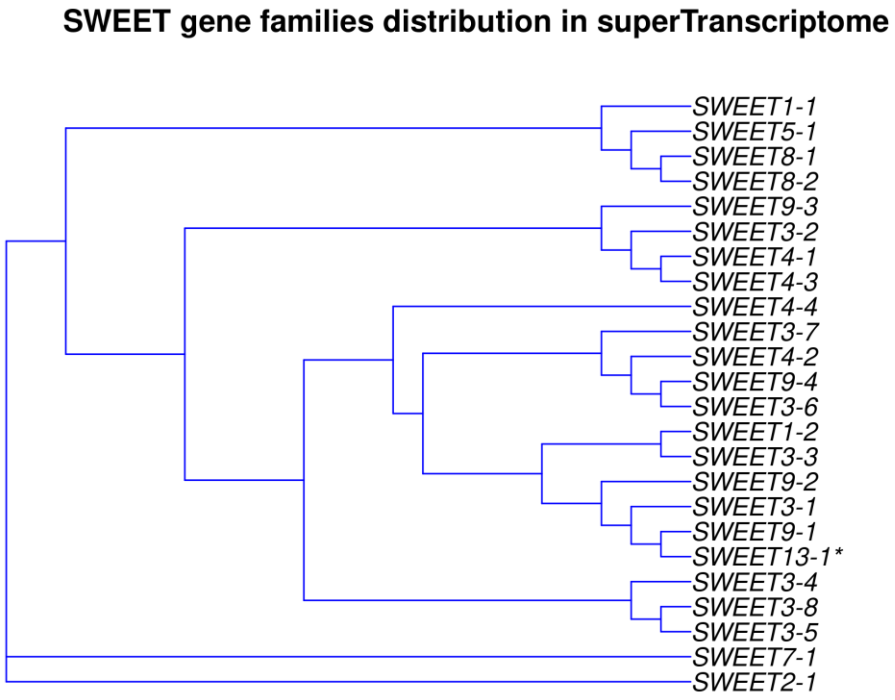


**Supplemental Figure S18:** Multiple Sequence Alignment showing SWEET gene families distribution in superTranscriptome

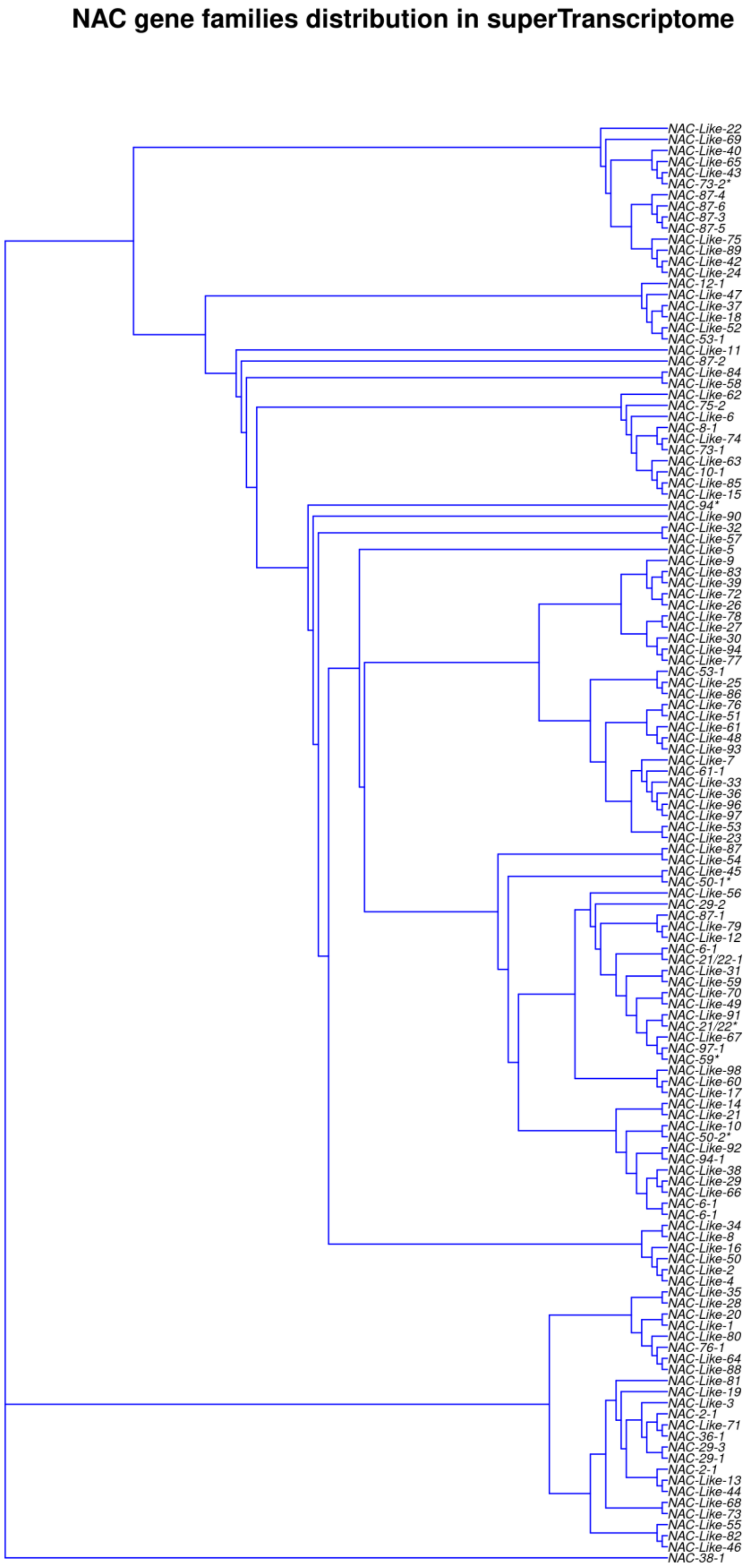


**Supplemental Figure S19:** Multiple Sequence Alignment showing NAC gene families distribution in superTranscriptome

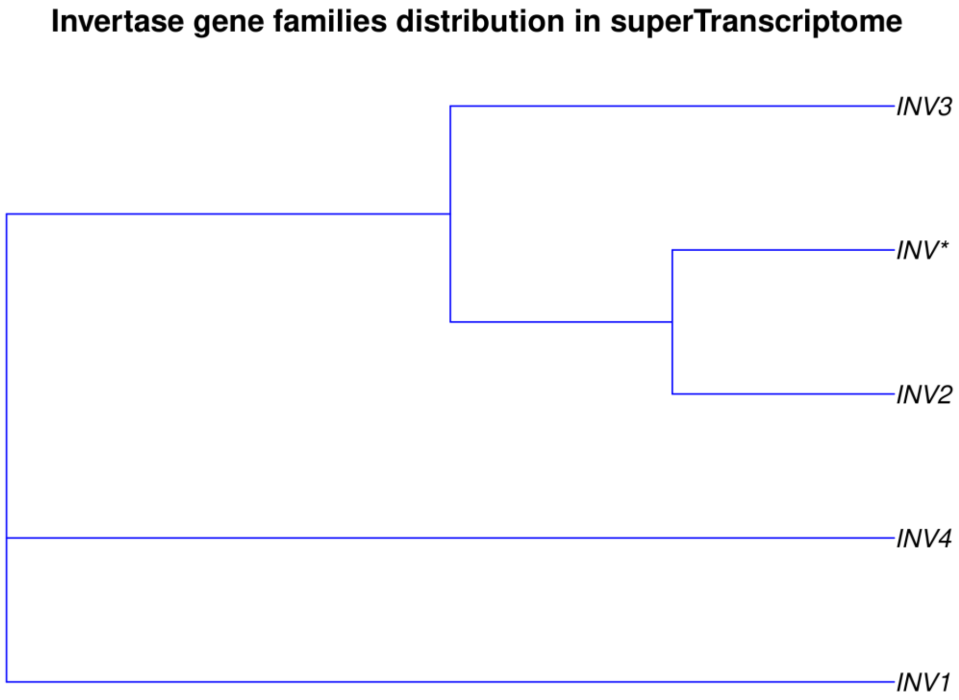


**Supplemental Figure S20:** Multiple Sequence Alignment showing Invertase gene families distribution in superTranscriptome

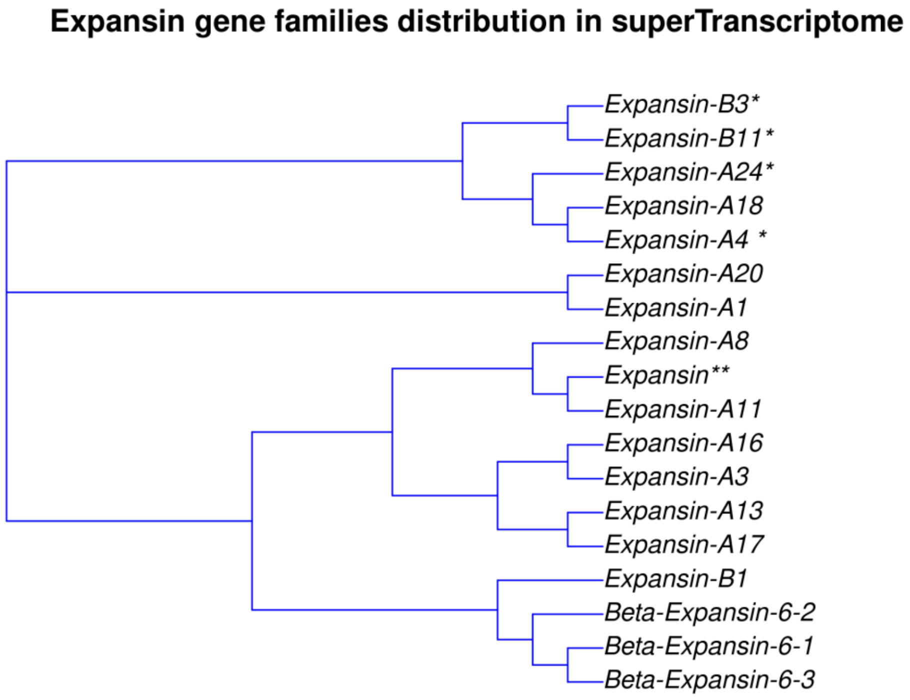


**Supplemental Figure S21:** Multiple Sequence Alignment showing Expansin gene families distribution in superTranscriptome

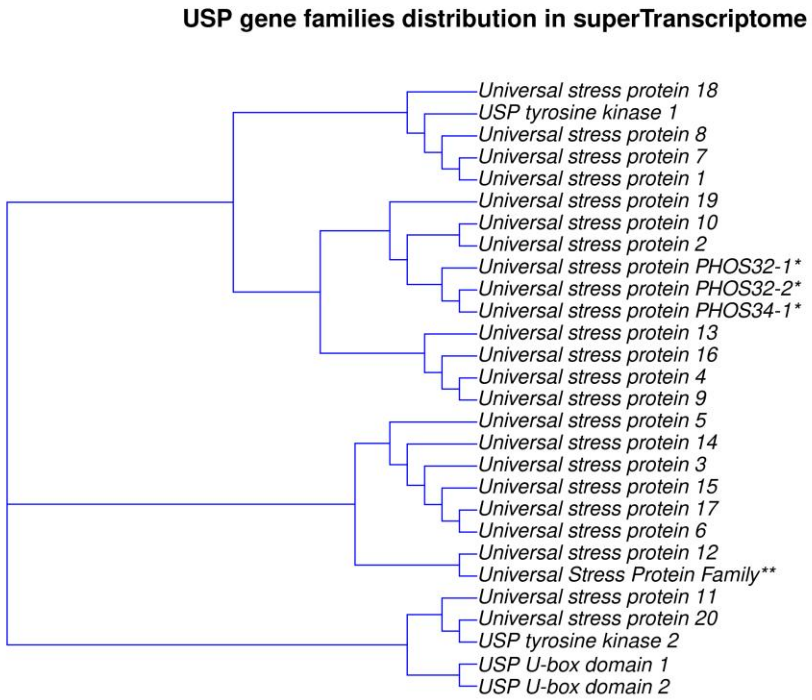


**Supplemental Figure S22:** Multiple Sequence Alignment showing USP gene families distribution in superTranscriptome

**Supplemental Table S1: Assembly statistics for superTranscriptome when clustered with different levels (70%, 80%, 90% and 98%) of sequence identity**

| Assembly Parameters | Description | superTranscriptome_70 | superTranscriptome_80 | superTranscriptome_90 | superTranscriptome_98 |
|---|---|---|---|---|---|
| n_seqs | the number of contigs in the assembly | 44719 | 44741 | 44826 | 45055 |
| Smallest | the size of the smallest contig | 96 | 96 | 96 | 96 |
| Largest | the size of the largest contig | 54581 | 54581 | 54581 | 54581 |
| n_bases | the number of bases included in the assembly | 99212176 | 100843832 | 100953715 | 102509949 |
| mean_len | the mean length of the contigs | 2217.83759 | 2253.21595 | 2251.39468 | 2274.49233 |
| n_under_200 | the number of contigs shorter than 200 bases | 204 | 204 | 204 | 204 |
| n_over_1k | the number of contigs greater than 1,000 bases long | 30245 | 30426 | 30466 | 30815 |
| n_over_10k | the number of contigs greater than 10,000 bases long | 454 | 454 | 454 | 456 |
| n_with_orf | the number of contigs that had an open reading frame | 29399 | 29434 | 29488 | 29764 |
| Mean_orf_percent | for contigs with an ORF, the mean % of the contig covered by the ORF | 49.4417 | 47.88405 | 47.8874 | 46.99359 |
| N90 | the largest contig size at which at least 90% of bases are contained in contigs at least this length | 1185 | 1203 | 1201 | 1218 |
| N70 | the largest contig size at which at least 70% of bases are contained in contigs at least this length | 2280 | 2333 | 2330 | 2359 |
| N50 | the largest contig size at which at least 50% of bases are contained in contigs at least this length | 3347 | 3408 | 3406 | 3432 |
| N30 | the largest contig size at which at least 30% of bases are contained in contigs at least this length | 4756 | 4805 | 4802 | 4809 |
| N10 | the largest contig size at which at least 10% of bases are contained | 8206 | 8191 | 8188 | 8150 |

| | | | | | |
|---|---|---|---|---|---|
| | in contigs at least this length | | | | |
| GC | % of bases that are G or C | 0.48767 | 0.48577 | 0.48578 | 0.48465 |
| Bases_n | the number of bases that are N | 84 | 84 | 84 | 80 |
| Proportion_n | the proportion of bases that are N | 0 | 0 | 0 | 0 |
